# The Role of Notch Signaling in Endometrial Mesenchymal Stromal/Stem-like Cells Maintenance

**DOI:** 10.1101/2022.02.09.479815

**Authors:** Sisi Zhang, Rachel W.S. Chan, Ernest H.Y. Ng, William S.B. Yeung

**Author notes:** **Co-correspondence:** Rachel W.S. Chan, Department of Obstetrics and Gynaecology, The University of Hong Kong, Pokfulam, Hong Kong; William S.B. Yeung, Department of Obstetrics and Gynaecology, LKS Faculty of Medicine, The University of Hong Kong, SAR, China. Equal contribution (Sisi Zhang, Rachel W.S. Chan).

## Abstract

Human endometrium undergoes cycles of regeneration in reproductive women. The endometrial mesenchymal stromal/stem cells (eMSC) contribute to this process. Notch signaling is essential for the homeostasis of somatic stem cells. However, its role in eMSC remains unclear. The gain and loss function shows that activation of Notch signaling promotes eMSC maintenance, while inhibition displays opposite effect. Activation of Notch pathway better maintains eMSC in a quiescent state. However, these quiescent eMSC can re-enter into the cell cycle depending on Notch and Wnt activity in the microenvironment, suggesting a crosstalk between two signaling pathways. In a mouse menstrual-like model, we observe that Notch signaling is highly involved in the dynamic endometrial remodeling event. Suppression of Notch signaling significantly reduces the proliferation of Notch1+ label-retaining stromal cells and consequently delays the endometrial repair. Our data demonstrate the importance of Notch signaling in regulating the endometrial stem/progenitor cells in vitro and in vivo.

## Introduction

Human endometrium undergoes cyclic proliferation, differentiation and shedding during women’s reproductive years (1). The regeneration of the endometrium after menses is an essential component of a menstrual cycle in humans and primates (2, 3). Endometrial stem/progenitor cells are vital in tissue regeneration after menstruation (4). Schwab and Gargett isolated a rare population of human endometrial mesenchymal stromal/stem-like cells (eMSC) based on co-expression of two perivascular markers: CD140b and CD146 (5). These cells showed somatic stem cell properties and phenotypic expression similar to other mesenchymal stem/stromal cells (6, 7).

Notch is a highly conserved signaling pathway that plays a vital role in tissue homeostasis and maintenance of somatic stem cells (8). Specific ligand-receptor interaction activates Notch signaling, leading to a series of proteolytic events and release of Notch intracellular domain (NICD). The cleaved NICD is then translocated to the nucleus where it provokes transcription of Notch target genes such as *HEY1, HEY2* and *HES1* (9). In human endometrium, several Notch family members are expressed across the menstrual cycle (10). Dysregulation of Notch1, DLL1 and JAG1 has been observed in endometrium from infertile women, suggesting an association of Notch signaling with infertility (11). Notch signaling has also been implicated in endometrial remodeling events such as decidualization and embryo implantation (10) and uterine-specific Notch1-knockout mice exhibit decidualization defect (12). Decreased Notch signaling is also associated with endometriosis (13). Gene expression profiling reveals increased activation of Notch signals in eMSC compared with endometrial stromal fibroblasts (eSF) (14). However, the role of Notch signals on proliferation and maintenance of eMSC remains largely undefined.

Another signaling pathway known to be involved in self-renewal and maintenance of somatic stem cells is WNT/β-catenin pathway (15). Myometrial cells can facilitate self-renewal of eMSC through the WNT5A/β-catenin signaling (16). Moreover, soluble secretory factors from endometrial niche cells at menstruation modulate the eMSC biological activities via activation of the WNT/β-catenin signaling (17). Accumulating evidence suggests a crosstalk between the Notch and the WNT/β-catenin signaling in somatic stem cells (18, 19). We postulate that such interaction exists in eMSC.

To explore the role of Notch signaling on the biological activities of endometrial stem/progenitor cells, we examined the expression of Notch activity in different subpopulations of human endometrial stromal cells and determined the interaction of Notch signaling with WNT/β-catenin signaling in eMSC *in vitro*. The functional significance of Notch signals in endometrial regeneration was studied in a mouse model of endometrial breakdown and regeneration with the use of label retaining cell (LRC) technique for identification of endometrial stem/progenitor cells *in vivo*.

## Results

### Activation of Notch signaling maintains phenotypic expression of eMSC

To determine whether Notch signaling plays a role on the maintenance of eMSC, we firstly compared the endogenous level of Notch target genes in eSF and eMSC. The relative mRNA levels of *HES-1* (Fig 1A, n=12, P<0.001) and *HEY-L* (Fig 1B, n=10, P<0.05) in eMSC were significantly higher than those in eSF. The relative mRNA levels of *HEY-1* (Fig 1C) and *HEY-2* (Fig 1D) tended to be higher in eMSC but the difference did not reach statistical significance (*HEY-1*: n=10, P=0.12; *HEY-2:* n=9, P=0.70).

**Figure 1.**
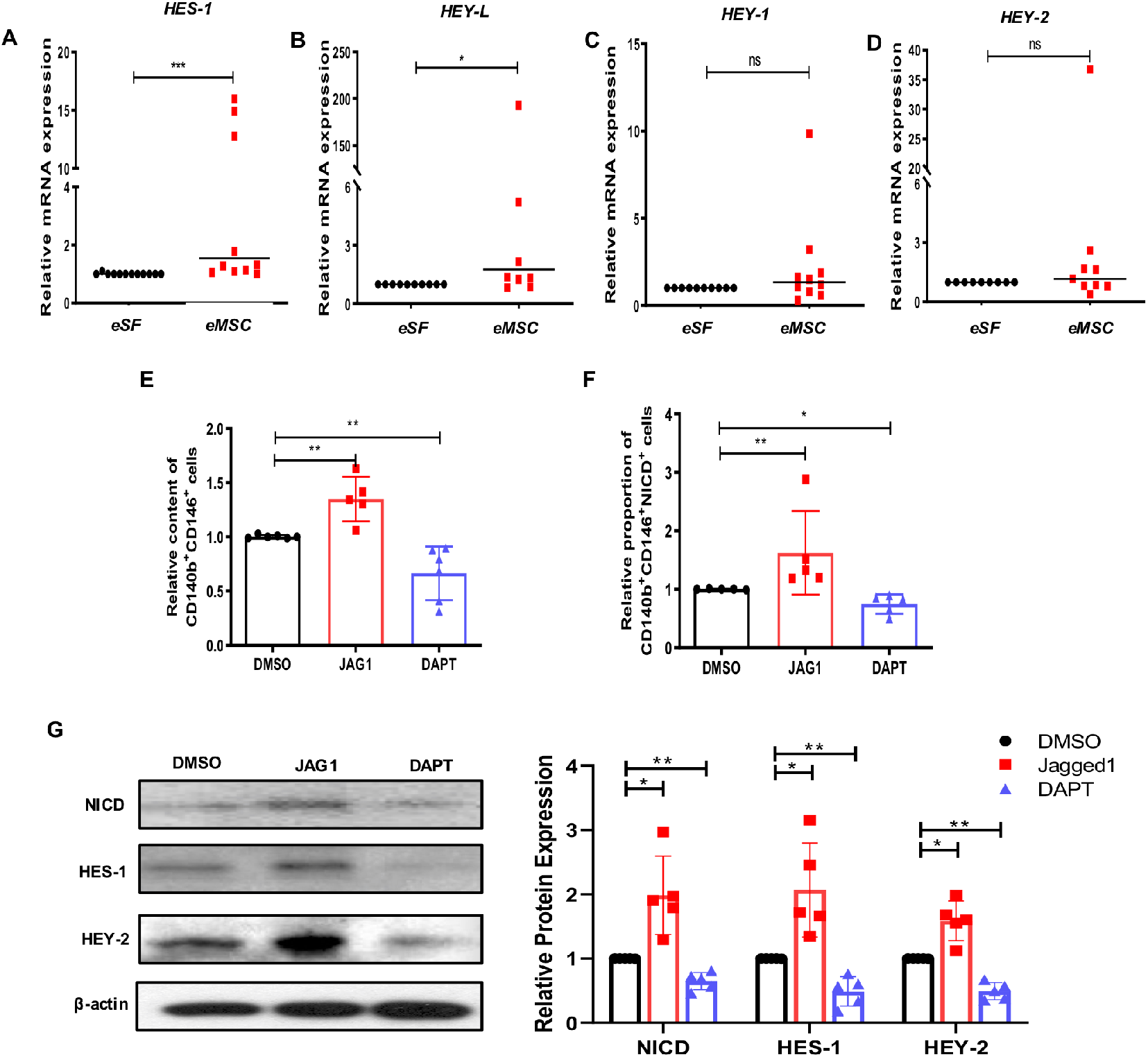
Activation of Notch signaling maintains phenotypic expression of eMSC. **(A-D)** The relative expression of Notch target gene in eSF and eMSC (n=9-12). **(E)** The relative content of CD140b^+^CD146^+^ cells after Notch signaling activation or inhibition by flow cytometry (n=6). **(F)** Quantitative analysis of CD140b^+^CD146^+^NICD^+^ cells after Notch signals activation or inhibition (n=5). **(G)** Representative western blotting images and quantitative analysis of Notch signals protein expression (n=5). Results are presented as mean ± SD; *P < .05; **P < .01; ***P < .001. Statistical analysis was performed using a two-tailed unpaired Student’s t test for parametric data and Mann-Whitney U test for non-parametric data, Kruskal-Wallis test followed by Dunn’s post-test for multiple group comparison. Abbreviation: eSF, endometrial stromal fibroblast; eMSC, endometrial mesenchymal stem-like cells, NICD, notch intracellular domain.

The gain and loss of function approaches were used to assess the role of Notch signaling on phenotypic expression of eMSC. The percentage of CD140b^+^CD146^+^ cells was higher when cultured on JAG1-coated plates than control group (Fig 1E, n=6, P<0.01) and treatment with DAPT to inhibit the γ-secretase complex abolished the difference (Fig 1E, n=6, P<0.01). Immunofluorescence staining showed increased nuclear accumulation of NICD (CD140b^+^CD146^+^NICD^+^ cells, Fig 1F, Fig S1, n=5, P<0.01) in eMSC cultured on the JAG1-coated plates, supporting activation of the Notch signaling. Concordantly, DAPT treatment reduced the proportion of NICD^+^ eMSC (Fig 1F, Fig S1, n=5, P<0.01).

The biological action of JAG1 on activation of the Notch signaling was confirmed by western blotting. JAG1 enhanced the protein expression of NICD, HES-1 and HEY-2 in eMSC compared to DMSO control (Fig 1G, n=5, P<0.05). The expression of these Notch target proteins was reduced after treatment with DAPT (Fig 1G, n=5, P<0.01).

### Notch1 mediates the maintenance effect of JAG1 on eMSC

Next, we evaluated the expression of Notch ligands and receptors in eSF and eMSC. Our results showed a non-significant higher expression level of *Notch1, Notch2* and *Notch3* mRNA in eMSC than in eSF (Fig S2A, *Notch1*: n=13, P=0.08; *Notch2*: n=10, P=0.12; *Notch3:* n=10, P=0.12). Interestingly, western blotting (Fig S2B, n=5, P<0.01) and immunofluorescence staining (Fig S2C, n=5, P<0.05) revealed significantly higher Notch1 protein expression in eMSC than in eSF. However, its expression level in eMSC were similar across the menstrual cycle (Fig S2D). Moreover, triple immunofluorescence staining confirmed the co-expression of CD140b, CD146 and Notch1 in 70% of the freshly isolated eMSC (Fig S2E).

The expression of JAG1, JAG2 and DLL4 in eMSC at different stage of the menstrual cycle was also investigated by western blotting (Fig S3A, S3B, n=3). Only JAG1 protein was consistently expressed in eMSC throughout the cycle (Fig S3A, S3B, n=3). The JAG1 protein expression in eSF and eMSC was similar (Fig S3C, n=5, P=0.68).

To study the action of JAG1–Notch1 on eMSC, we evaluated the binding of fluorescence labeled JAG-1 after knockdown of Notch1. The fluorescence intensity of bound JAG1 on eMSC was reduced after transfection with Notch1 siRNA (Fig S2G, n=6, P<0.01). Knockdown of Notch1 also abolished the maintenance effect of JAG1 on eMSC phenotypic expression (Fig S2H, n=7, P<0.01).

### Activation of Notch signaling maintains eMSC in a quiescent state

Notch signaling is required for maintenance of quiescent somatic stem cells (20, 21). Therefore, we assessed the proliferation activity of eMSC after activation and inhibition of Notch pathway. The rate of proliferation in eMSC was significantly lower after culture on JAG1 coated plate than on uncoated plate (Fig 2A, n=7, P<0.05), while no difference was detected between the DAPT treated and control group (Fig 2A, n=7, P=0.68). The mRNA and protein expression of the proliferation marker Ki67 in eMSC were also lower after culture on JAG1 coated plate than uncoated plate (Fig 2B, 2C, n = 6, P<0.05). We also evaluated the confluence of eMSC after culture for 7 days. Brightfield images collected during kinetic experiment demonstrated the eMSC were not in direct contact suggesting that the decrease in proliferation was not related to contact inhibited cultures (Fig S4A).

**Figure 2.**
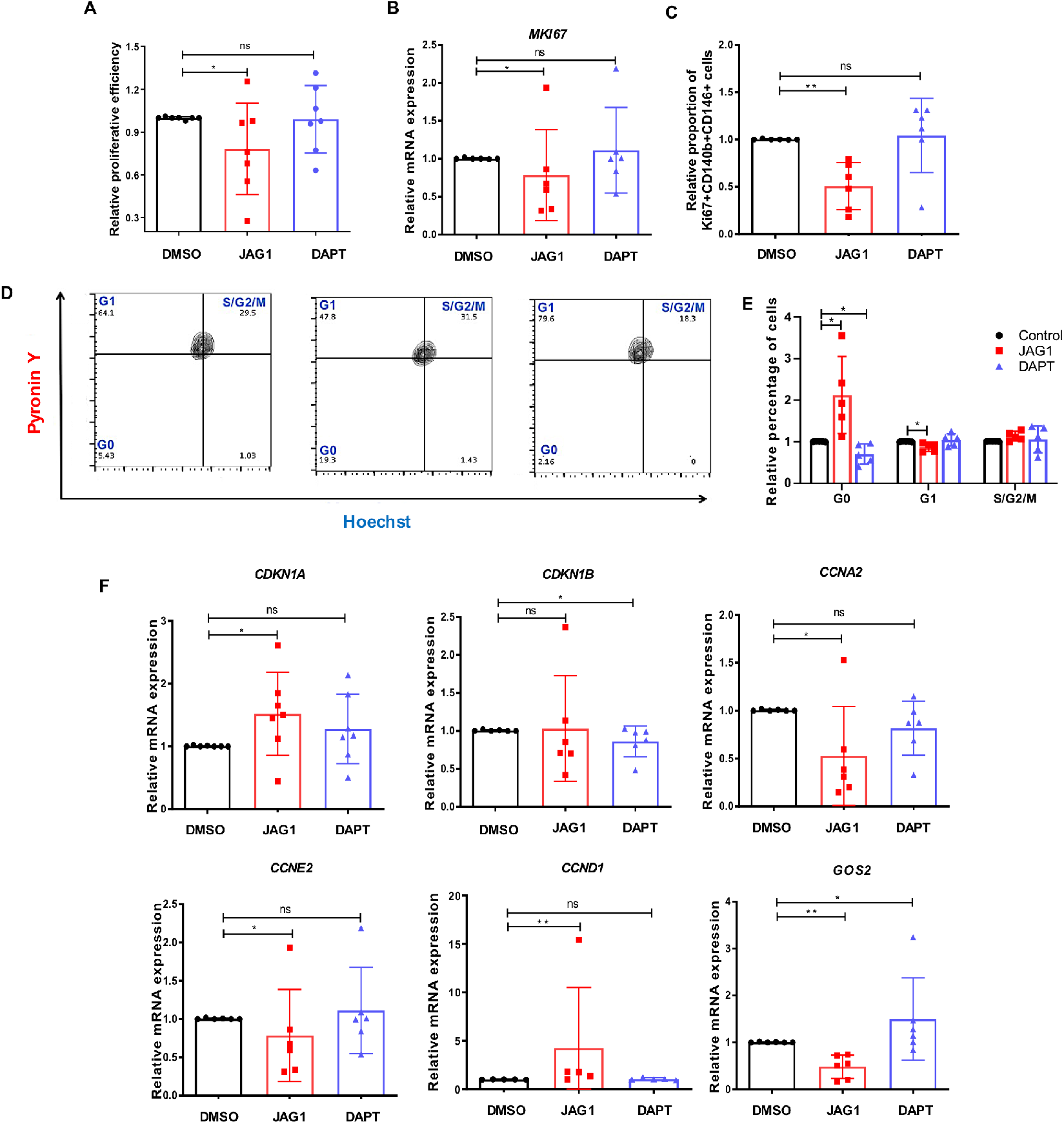
The role of Notch signaling pathway on maintaining eMSC at quiescent state. **(A)** The relative proliferative efficiency by proliferation kit (n=7). **(B)** The relative gene expression of MI67 after Notch signals activation or inhibition (n=6). **(C)** The relative percentage of CD140b^+^CD146^+^Ki67^+^ cells after Notch signals activation or inhibition by flow cytometry (n=6). **(D)** Representative images of Hoechst and Pyronin Y staining on eMSC by flow cytometry (n=5). **(E)** The relative proportion of eMSCs on different cell cycle by Hoechst and Pyronin Y staining (n=5). **(F)** The relative cell cycle related gene expression on eMSC after Notch signals activation or inhibition (n=5-7). Results are presented as mean ± SD; *P < .05; **P < .01. Statistical analysis was performed using a Kruskal-Wallis test followed by Dunn’s post-test. Abbreviation: eMSC, endometrial mesenchymal stem-like cells.

To investigate whether the reduction in proliferation activity is related to quiescence of stem cells, we analyzed the cell cycle status of eMSC. Fig 2D shows the cell cycling status of eMSC with Notch activation and inhibition by flow cytometry analysis. JAG1 induced more eMSC at the G0 state and decreased the proportion of cells in the G1 state (Fig 2E, n=5, P<0.05). Blocking the Notch signaling by DAPT reversed the effect (Fig 2E, n=5, P<0.05). Moreover, DAPT at the concentration used in this study did not induce apoptosis as the treatment had no effect on the mRNA expression of the proapoptotic *BCL-2* and the antiapoptotic *BAX*, though the treatment reduced that of the inhibitor of apoptosis gene *BIRC5* (Fig S4B-4D).

Consistent with the cell cycle analysis, activation of the Notch signaling pathway upregulated the mRNA expression of *CDKN1A* (Fig 2F, n=7, P<0.01), *CCND1* (Fig 2F, n=5, P<0.05) and downregulated that of *CCNA2* (Fig 2F, n=6, P<0.05), *CCNE2* (Fig 2F, n=6, P<0.05) and *GOS2* (Fig 2F, n=6, P<0.01). Inhibition of Notch activity with DAPT reduced the expression of *CCKN1B* (Fig 2F, n=6, P<0.05) but elevated that of *GOS2* (Fig 2F, n=6, P<0.05). These findings suggest activation of Notch signaling can better maintain eMSC in a quiescent state.

### WNT signaling activates the quiescent eMSC

To determine if the JAG1-induced quiescent state of eMSC was reversible, we examined the proliferation activity after removal of JAG1 in the subsequent cell passage. At passage 0 (P0), eMSC were cultured on JAG1 coated plates and in the following cell passages (P1 - P2) the cells were cultured on uncoated plate. JAG1 treatment decreased the proliferation of eMSC at P0 (Fig 2A). However, the phenomenon was not observed in eMSC at P1 (Fig 3A, n=9, P=0.95) and P2 (Fig 3B, n=6, P=0.16). To our surprise, the clonogenicity of eMSC was lower when JAG1 was depleted at P1 than the control (Fig 3C, 3D, n=9, P<0.05). The difference was not observed at P2 (Fig 3C, 3D, n=9, P=0.62).

**Figure 3.**
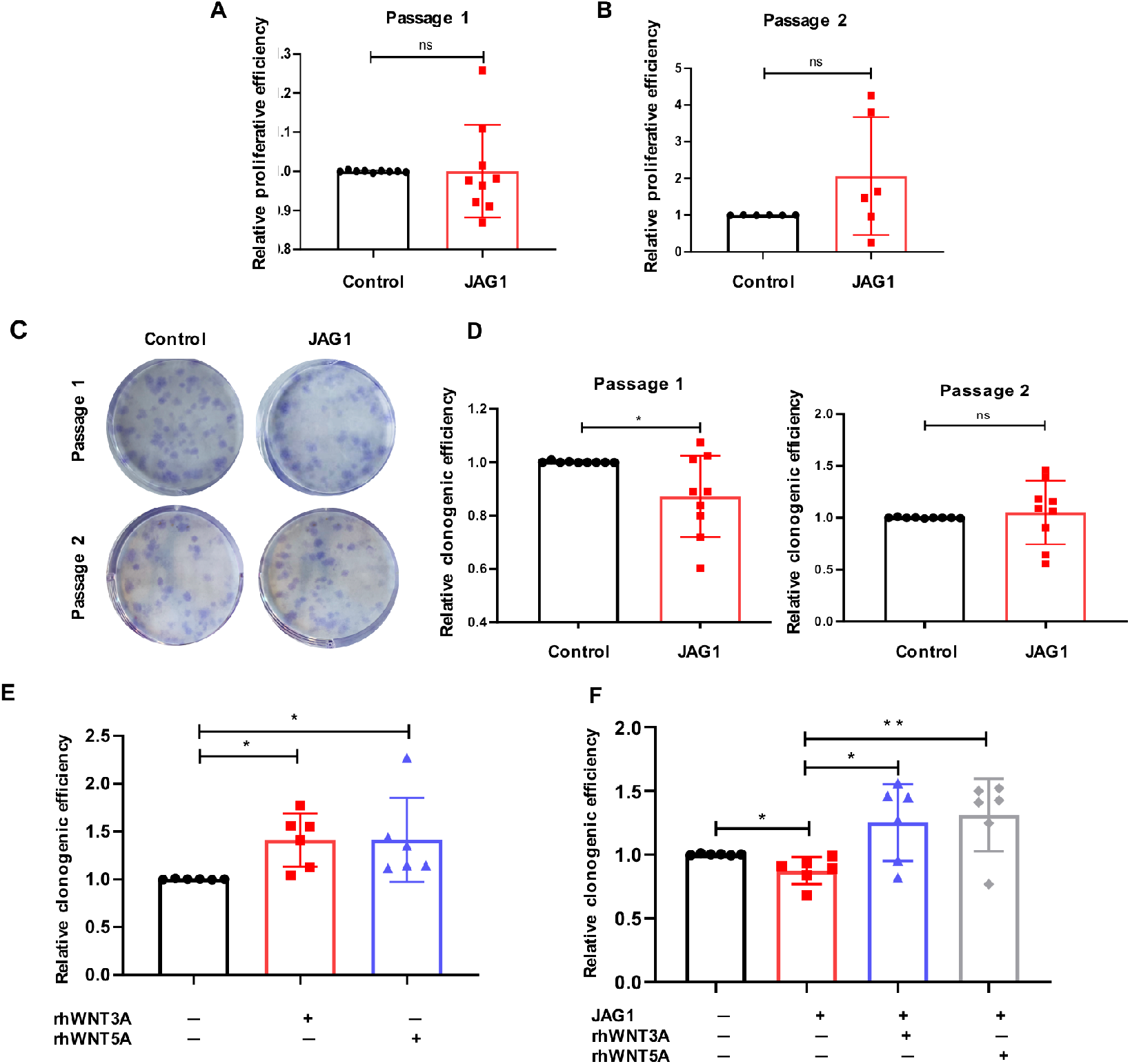
Notch activation can induce reversible arrest in eMSC. The relative proliferative efficiency of eMSC at **(A)** passage 1 and **(B)** passage 2 after cultured with JAG1 at passage 0 (n=6-9). **(C)** Representative image showing the total colony formation of eMSC at passage 1 and passage 2 after cultured with JAG1 at passage 0. **(D)** The relative clonogenic activity of eMSC at passage 1 and passage 2 after cultured with JAG1 at passage 0 (n=9). **(E)** The relative clonogenic activity of eMSC under different conditions (n=6). **(F)** The relative clonogenic activity of eMSC under different conditions after cultured with JAG1 at passage 0 (n=6). Results are presented as mean ± SD; *P < .05; **P < .01. Statistical analysis was performed using a two-tailed unpaired Student’s t test for two group comparison, Kruskal-Wallis test followed by Dunn’s post-test for multiple group comparison. Abbreviation: eMSC, endometrial mesenchymal stem-like cells; rh, recombinant.

Wnt ligands promote the formation of eMSC colonies(16) (Fig 3E, n=6, P<0.05). Thus, we hypothesis that Wnt3A or Wnt5A may activate the quiescent eMSC. The action of JAG1 in the presence of Wnt3A or Wnt5A on colony formation was evaluated. Indeed, the inhibitive effect of JAG1 on the clonogenicity of eMSC at P1 reversed upon treatment with Wnt3A or Wnt5A (Fig 3F, n=6, P<0.05). Taken together, these findings suggest the JAG1-induced quiescent eMSC could re-enter into the cell cycle upon Wnt activation.

### Interaction of NOTCH and WNT/β-catenin pathway in eMSC

#### Action of NOTCH activity on active β-catenin

Crosstalk between WNT and Notch pathway has been reported in different systems (19). Therefore, we investigated the existence of similar phenomenon in eMSC. Immunofluorescent staining showed that the expression of active β-catenin in eMSC increased after activation and suppressed upon inhibition of Notch signaling (Fig S5, Fig 4A, n=5, P<0.01). The TCF/LEF transcriptional activity of eMSC shown in the reporter assay confirmed the observation (Fig 4B, n=5, P<0.01). The protein expression of active β-catenin increased significantly upon activation and decreased after inhibition of the Notch pathway (Fig 4C, n=5, P<0.01). As expected, PLA revealed that eMSC cultured on JAG1 coated plate had more binding between NICD and active β-catenin in the nuclear region than the DMSO control. In contrast, the nuclear PLA positive signal reduced after treatment with DAPT (Fig 4D, n=3).

**Figure 4.**
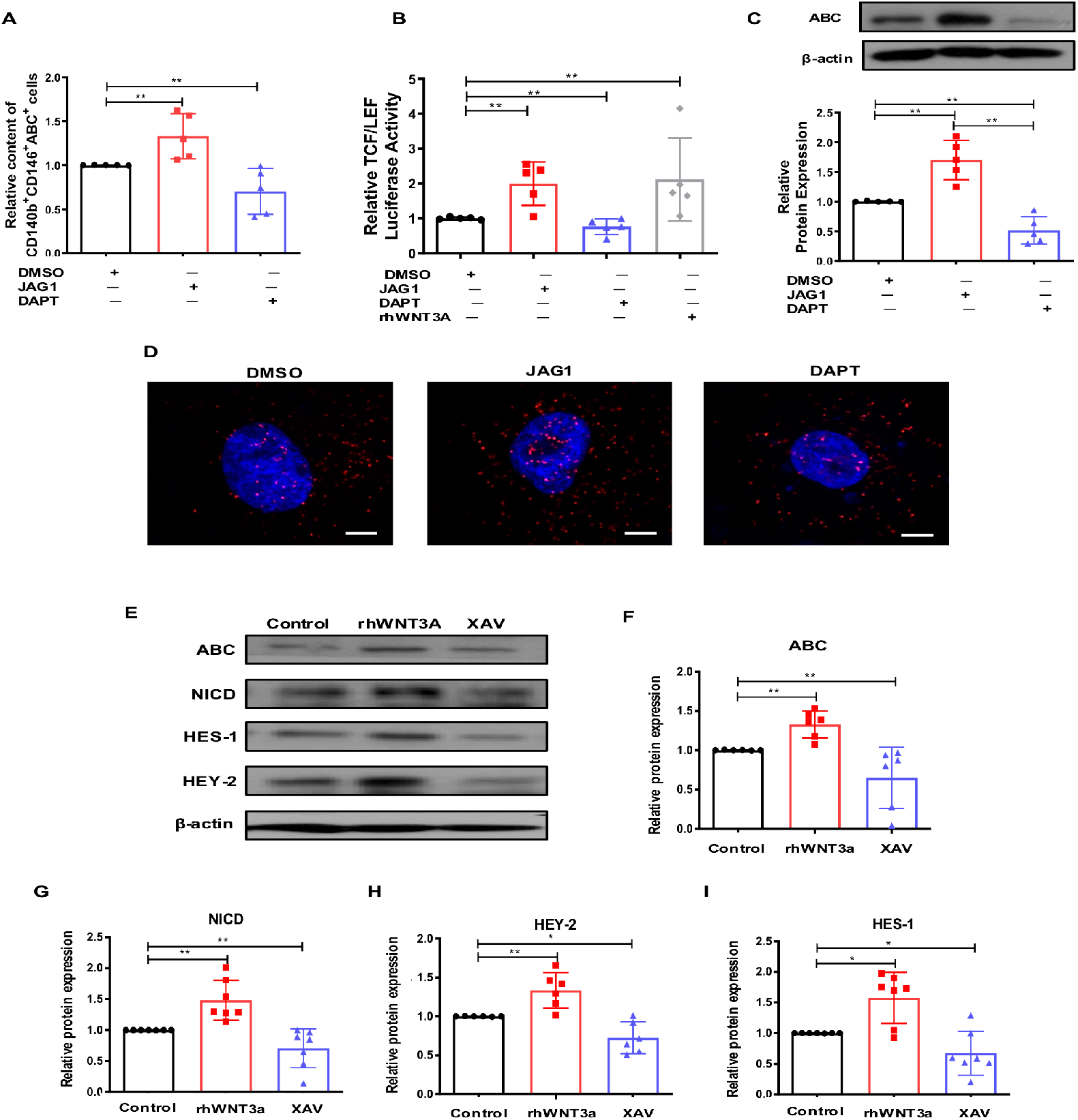
Interaction of Notch and WNT/β-catenin pathway in eMSC. **(A)** Quantitative analysis of CD146^+^CD140b^+^ABC^+^ endometrial stromal cells after Notch signals activation or inhibition (n=5). **(B)** The TCF/LEF luciferase signal of eMSC after Notch signals activation or inhibition (n=5). **(C)** Representative western blotting images and quantitative analysis of ABC protein expression on eMSC after Notch signals activation or inhibition (n=5). **(D)** Representative images of *in situ* proximity ligation assay on eMSC (n=3), scale bar: 10 µm. **(E)** Representative western blotting images of active β-catenin (ABC) and Notch signals protein expression in eMSC after canonical WNT signaling activation or inhibition (n = 6-7). **(F-I)** Quantitative analysis of active β-catenin and Notch signals protein expression in eMSC after canonical WNT signaling activation or inhibition (n= 6-7). Results are presented as mean ± SD; *P < .05; **P < .01. Statistical analysis was performed using a Kruskal-Wallis test followed by Dunn’s post-test. Abbreviation: eMSC, endometrial mesenchymal stem-like cells; ABC, active β-catenin; rh, recombinant.

#### Action of WNT activity on NOTCH related proteins

Next, we investigated the functional role of canonical WNT signals on the expression of Notch related proteins in eMSC. Recombinant WNT3A and XAV939 were used to activate or inhibit the WNT signals, respectively (Fig 4E, 4F, n=6, P<0.01). Recombinant WNT3A remarkably increased the expression of Notch related proteins NICD, HEY-2 and HES-1 in eMSC (Fig 4E, 4G-4I, n=6-7, NICD & HEY-2: P<0.01; HES-1: P<0.05). In contrast, the WNT inhibitor XAV939 downregulated the Notch downstream activities (Fig 4E, 4G-4I, n=6 - 7, NICD: P<0.01, HEY-2 & HES-1: P<0.05).

### Endometrial label retaining stromal cells (LRSC) in a mouse menstrual-like model

A mouse menstrual-like model was employed to gain insight into the role of Notch activity on stem cells during endometrial regeneration *in vivo* (22). The label retaining cells approach was used to identify the endometrial stem/progenitor in animals because of absence of defined stem cells markers for mouse endometrium. In the mouse model, sesame oil was injected into one uterine horn of the mice on day 4 of pseudopregnancy to induce decidualization, which resulted in increase of the size and obliteration of the lumen of the treated horn on day 7. Enlargement of the treated horn continued to day 9. During this time course, the color of the horn changed from pink to dark red/purple, indicating readiness of tissue breakdown. The size of decidualized uterine horn decreased on day 10 and became macroscopically the same as the control on day 12 (Fig S6B).

Histological examination showed an open lumen and intact endometrium in the uterine horn before decidualization. Three days after oil injection (day 7), typical features of decidualization including intense vascularization, densely packed decidualized stromal cells and closure of the lumen were observed. Tissue breakdown occurred on day 9, resulting in slough off the decidualized tissue into the lumen. Repair of the endometrium commenced on day 10 when the luminal epithelium re-epithelialized and the stromal cells began to proliferate. The morphology of the endometrium resembled the control, and the repair was completed by day 12 (Fig S6C).

The stem/progenitor cells of mice were labeled with BrdU in prepubertal state. We showed previously a steady percentage of BrdU^+^ cells after a 6-week chase (23). Hence, the BrdU^+^ cells after a 6-week chase were referred as LRSC in the present menstrual-like model, and they accounted for 2.73 ± 0.33% (n=4) of the total stromal cells (Fig 5A, 5B). At decidualization, the percentage of BrdU^+^ stromal cells showed a rapid decline compared to that of the control (Fig 5A, 5B, n=4, P<0.001). Only 0.38 ± 0.05% (n=4) of LRSC were detected in the decidualized uterine horn (Fig 5A, 5E). The number of LRSC increased at endometrial breakdown (Fig 5A, 5B, Day 9: 1.32 ± 0.14%, n=5, P<0.001) but began to decline at early and late repair (Fig 5B, Day 10: 1.15 ± 0.23% n=4; Day 12: 0.68 ± 0.13%, n=3, P < 0.05). Overall, the percentages of LRSC in the decidualized uteri horn during repair were lower than the control (Fig 5B, early repair: P<0.05, n=4; late repair: P<0.01, n=3).

**Figure 5.**
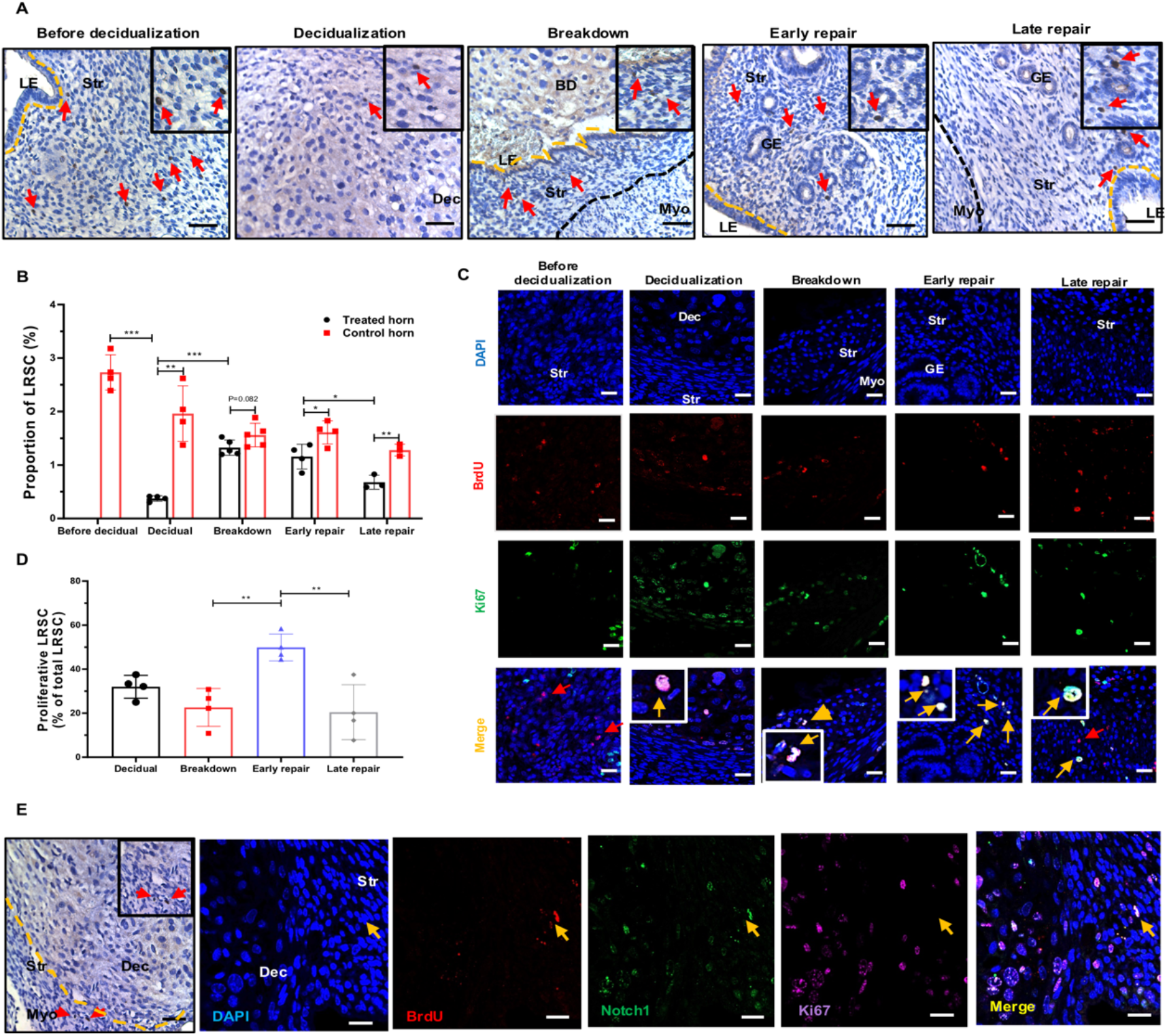
Localization of BrdU-labeled stromal cells in mouse menstrual-like model. **(A)** Localization of LRSC on endometrium in mouse menstrual-like model. *Inserts* are enlarged figures of BrdU-labeled cells. Red arrows show the distribution of LRSC at different time points. **(B)** BrdU-labeled stromal cells are expressed as a percentage of total stromal cells. **(C)** Representative immunofluorescent images show LRSC colocalizing with proliferating marker Ki67 (yellow arrow) at decidualization, breakdown, early and late repair. No colocalization with Ki67 (red arrow) for LRSC before decidualization. **(D)** The percentage of proliferating LRSC in mouse menstrual-like model. **(E)** LRSC remained in the dense stromal compartment during decidualization. Yellow arrows indicate no localization of Ki67 on LRSC. *Inserts* are enlarged figures of BrdU-labeled cells, scale bar: 20 µm. Results are presented as mean ± SD; *P < .05; **P < .01; ***P < .001. n = 3–5 per group. Statistical analysis was performed using a two-tailed unpaired Student’s t test for two group comparison, Kruskal-Wallis test followed by Dunn’s post-test for multiple group comparison. Abbreviation: BrdU, bromodeoxyuridine; BD, breakdown; Dec, decidua; GE, glandular epithelium; LE, luminal epithelium; LRSC, label retaining stromal cells; Myo, myometrium; Str, stroma.

### Endometrial LRSC proliferate during repair

Proliferation of LRSC was evaluation by co-expression of BrdU and Ki67. No proliferating LRSC were found in the endometrium before decidualization, suggesting that these cells remained in a quiescent state (Fig 5C, 5D). A high percentage of proliferating LRSC was observed (31.99 ± 5.20%, n=4) in decidualized tissue. The percentages remained high until tissue breakdown (22.62 ± 8.61%, n=4). The percentage of proliferating LRSC was highest at early repair and declined rapidly at late repair (Fig 5C, 5D, early repair: 49.86 ± 6.09%, n=4; late repair: 20.46 ± 12.49%, n=4, P<0.01), which might contribute to the decrease of LRSC observed at late repair (Fig 5B). Interestingly, a small population of LRSC near the endometrial/myometrial junction did not express Ki67 during decidualization (Fig 5E). We postulate these non-proliferating LRSC will be functional during endometrial remodeling (Fig 5D).

### Proliferating endometrial LRSC express Notch1

The expression of Notch1 in LRSC during endometrial remodeling was investigated. Before decidualization, 25.0 ± 5.0% of the LRSC were Notch1^+^. The percentage increased to 44.88 ± 44.37% (Fig S7A, S7B, n=4, P<0.05) in decidualized endometrium. The percentage of Notch1^+^ LRSC remained relatively constant until late repair when it decreased to 30.87 ± 5.57%, which was significantly lower than that in early repair (Fig S7A, S7B, n=4, P<0.05). Interestingly, Notch1^+^ LRSC proliferated (BrdU^+^Notch1^+^Ki67^+^ cells) only during events of endometrial remodeling i.e., at decidualization (14.24 ± 5.0%), tissue breakdown (9.16 ± 2.20%) and early repair (21.39 ± 17.42%, Fig S7C, S7D, n=3-5). Moreover, the mouse endometrium expressed Notch ligands JAG1 and DLL4 throughout these remodeling events (Fig S6D, S6E, n=3-5).

### DAPT treatment prolongs endometrial repair after menstrual-like breakdown

To determine the function of Notch signaling on LRSC, DAPT was delivered into the endometrial microenvironment on the day of breakdown. Blockage of Notch signals in the mouse endometrium was confirmed by western blotting, showing reduced expression of Notch1 and HES-1 expression after DAPT treatment (Fig S8A-8C, n=3, P<0.05).

In the vehicle treated group, the stromal compartment was fully restored, and re-epithelialization of the luminal epithelium with only a small portion of unrepair luminal epithelium was observed by early repair (Fig 6A). Endometrial regeneration was completed, and the luminal epithelium became intact by late repair. When DAPT was administrated into the uterine cavity, most part of the luminal epithelium was absent during early repair (Fig 6A). Even at late repair, sites without luminal epithelium remained, indicating incomplete regeneration of endometrium. Consistent with the morphologic changes, the endometrial thickness of the DAPT treated group was reduced remarkably compared with the vehicle group (Fig 6B, n=5, P<0.05). Taken together, these results suggest Notch signaling is required for post-menstrual regeneration and inhibition of this pathway delayed tissue repair. Moreover, the proportion of proliferating LRSC declined significantly in the DAPT treated group, which resulted in a higher proportion of LRSC detected at both early and late repair than vehicle group (Fig 6C, 6D, 7A, 7B, n=5, early repair: P<0.01, late repair: P<0.05).

**Figure 6.**
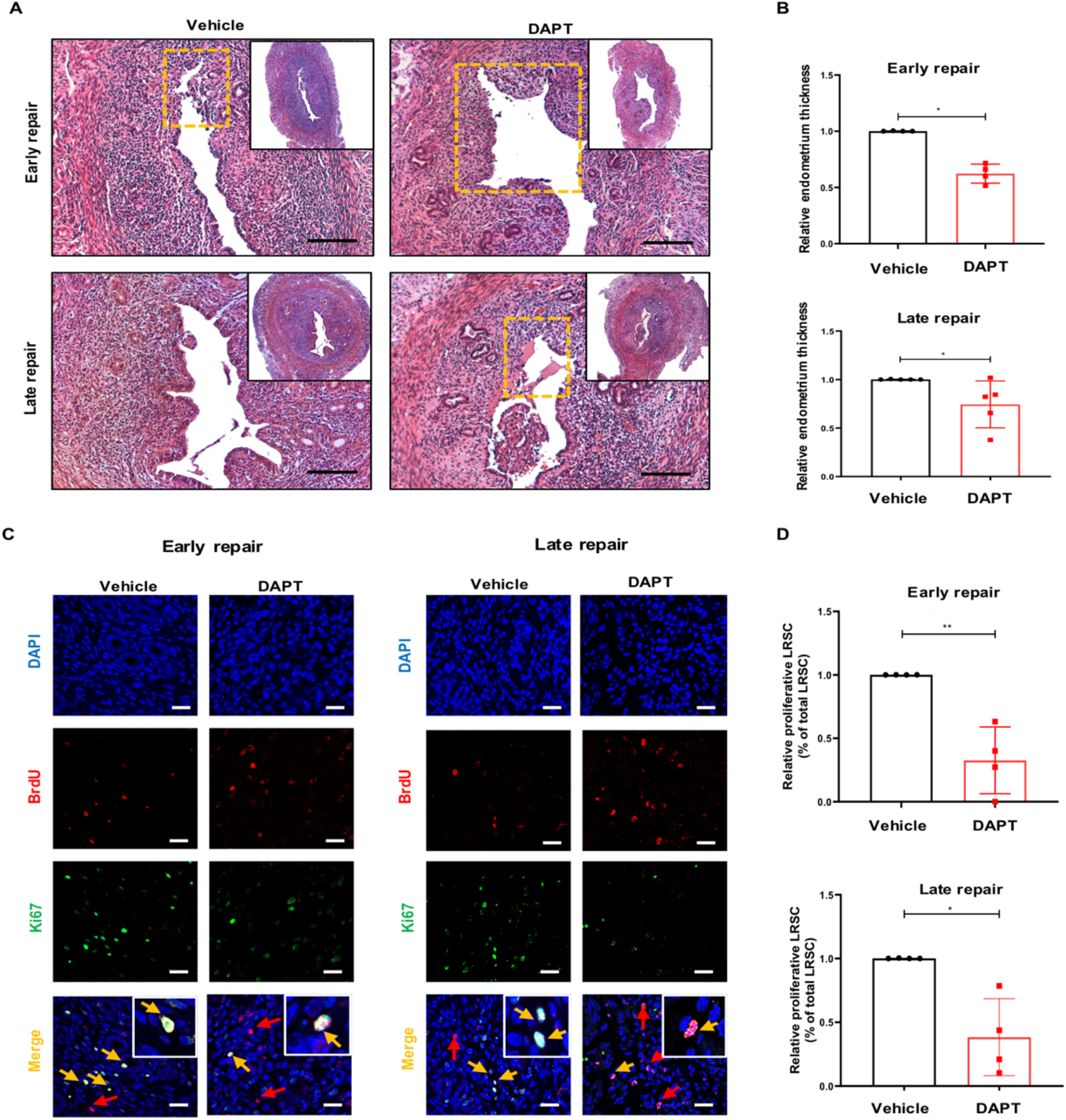
Proliferation of LRSCs after DAPT treatment. **(A)** Microscopic analysis of H&E-stained uteri after DAPT treatment at early and late repair. *Inserts* show the whole transverse section of the uterine tissue. Dashed boxes indicate unrepair sites. Scale bar: 50 µm. **(B)** Relative endometrium thickness after DAPT treatment at early and late repair. **(C)** Representative immunofluorescent images of proliferating LRSC after DAPT treatment at different time points. Yellow arrows show colocalization of proliferating marker Ki67 on LRSC. Red arrows indicate no localization of Ki67 on LRSC. **(D)** Relative percentage of proliferating LRSC after DAPT treatment at different time points. Scale bar: 20 µm. Results are presented as mean ± SD; *P < .05; **P < .01. n = 3–5 per group. Statistical analysis was performed using a two-tailed unpaired Student’s t test. Abbreviation: BrdU, bromodeoxyuridine; GE, glandular epithelium; LE, luminal epithelium; Myo, myometrium; Str, stroma.

Dual immunofluorescence staining revealed that the proportion of Notch1^+^ LRSC at early repair was lower in the DAPT treated group than the vehicle group (Fig 7C, 7D, n=5, P<0.001). The inhibitory effect continued at late repair (P<0.01). There were also less Notch1^+^ LRSC recruited into proliferation during early repair when treated with DAPT (Fig S8D-8F, n=3, P<0.01). The finding further explains the presence of more LRSC in the DAPT treated mice (Fig 7A, 7B).

**Figure 7.**
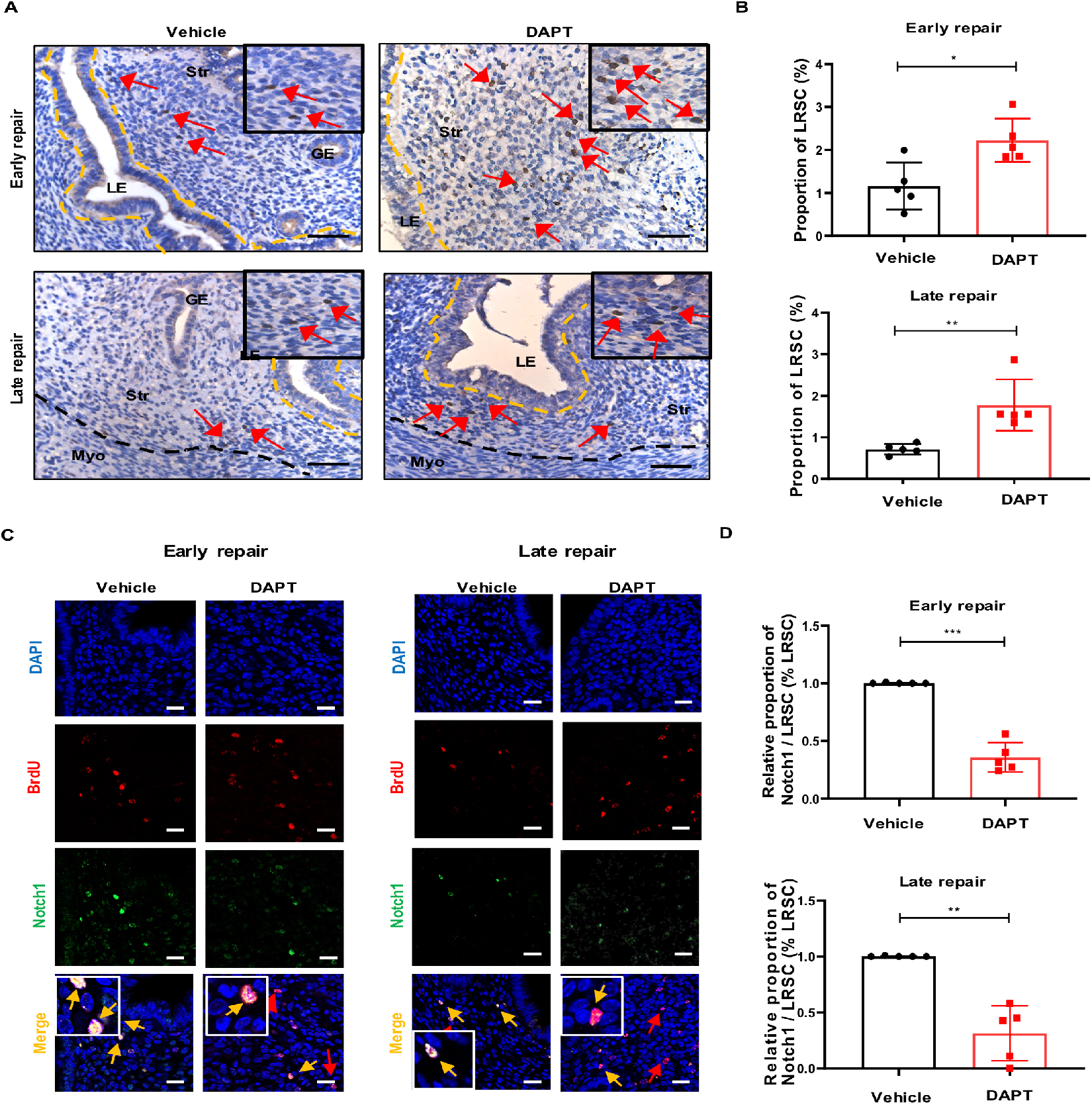
DAPT treatment increased the proportion of LRSC at early and late repair. **(A)** Localization of LRSC on uterine tissues after DAPT treatment at different time points. *Inserts* are enlarged figures of BrdU-labeled cells. Red arrows show the distribution of LRSC. **(B)** Proportion of LRSC after DAPT treatment at different time points. **(C)** Representative immunofluorescent images show LRSC colocalizing with Notch1 (yellow arrow) after DAPT treatment at early and late repair. Red arrows indicate no colocalization of Notch1 on LRSC. *Inserts* are enlarged figures of LRSC co-expressing Notch1. **(D)** Relative percentage of Notch1/ LRSC after DAPT treatment at different time points. Results are presented as mean ± SD; *P < .05; **P < .01. n = 3–5 per group. Scale bar: 20 µm. Statistical analysis was performed using a two-tailed unpaired Student’s t test. Abbreviation: BrdU, bromodeoxyuridine; GE, glandular epithelium; LE, luminal epithelium; Myo, myometrium; Str, stroma.

## Discussion

Since the discovery of eMSC in human endometrium, very few studies have sought to investigate their functional roles in endometrial repair after menstruation (1, 24). The underling mechanism that regulates the activities of eMSC remains largely unknown. In this study, we demonstrated that Notch signaling pathway plays a role in the maintenance of eMSC. Importantly, inhibition of Notch signals affected the function of stem/progenitor cells to repair after tissue breakdown in the mouse endometrium. To our knowledge, this is the first study exploring the potential role of Notch signals on endometrial stem/progenitor cells both *in vitro* and *in vivo*.

Many studies have reported Notch signaling is essential for homeostasis of somatic stem cells (8, 25). The high level of endogenous Notch related genes expressed by eMSC when compared to eSF suggest the involvement of Notch signaling pathway in stem cell regulation. This is consistent with the finding reported by Spitzer *et al*., where gene expression profiling showed an upregulation of Notch signals in eMSC (14). By manipulating the Notch signals with pharmacological agonist or antagonist, our results showed promotion of eMSC maintenance upon Notch stimulation or the reversed effect upon inhibition. Similar finding was detected in SUSD2^+^ eMSC (26). Together, these observations highlight a key role for Notch signals in the maintenance of endometrial stem cells.

The Notch family receptors are highly involved with stem cell maintenance and cell fate specification (25, 27). Ables *et al*. demonstrated that Notch1 is required for the maintenance of mice adult neural stem and progenitor cells (28). In bone marrow stromal/stem cells (BMSCs), Notch2 enhanced BMSCs self-renewal and proliferation *ex vivo* while their osteogenic and chondrogenic differentiation potential were preserved (29). In human endometrium, our results revealed that eMSC express Notch1. Knockdown of Notch1 receptor reversed the stimulatory effect of JAG1 on the activity of eMSC, demonstrating that eMSC required Notch1 for stem cell functions. Endothelial cells exhibit strong expression of JAG1 (14, 30). Since eMSC reside in perivascular regions across the human endometrium, it will be worth investigating the ligand-receptor interaction between endothelial cells and eMSC. We speculate Notch signals in eMSC will be activated with the adjacent endothelial cells to promote stem cell activities.

Quiescence and self-renewal are critical for reservation of somatic stem cells (31). Stem cells maintain in a quiescent state to protect the cells against the loss of self-renewal and the rapid exhaustion of stem cell pool (32). Here we demonstrated that activation of Notch signals by JAG1 prevented proliferation of eMSC and regulated the entrance of eMSC into the cell cycle. Moreover, our data showed that the JAG1 induced quiescent state on eMSC was reversible. These results are in agreement with other studies, where Notch signaling is a critical factor regulating quiescence in muscle and neural stem cells (33, 34). We propose that specialized signals from the niche cells during menstruation can stimulate eMSC to re-enter the cell cycle and execute physiological functions to restore the endometrial lining. While all notch ligand family members can be detected in the human endometrium. Only JAG1 and DLL1 were tightly regulated in the endometrium during the menstrual cycle (11). In future studies, it will be important to understand the action of these ligands on eMSC in controlling the balance between quiescent and activated state during the menstrual cycle.

Several studies have demonstrated the functional crosstalk between WNT and Notch signaling in somatic stem cells (35, 36). WNT signaling together with Notch signaling were required to regulate stemness and differentiation of human fallopian tube organoids (37). In human neural progenitor cells, promoting effect of WNT activation on neurogenesis was based on regulation of Notch-target genes (35). We previously demonstrated that myometrial cells regulated the self-renewal of eMSC via WNT/β-catenin signaling (16). Our current results showed that activation of WNT signaling promoted the quiescent eMSC to proliferate. These phenomena prompted us to investigate whether a crosstalk exist between Notch and WNT signaling in the modulation of eMSC. Indeed, further investigation revealed that inhibition of Notch signaling in eMSC can lead to the downregulation of active β-catenin. While activation of Notch signaling upregulated WNT activity in eMSC. Similar results were observed on the activity of Notch signals when WNT signaling was pharmacologically manipulated in eMSC. Taken together, our results suggest the interplay between Notch and WNT signaling play a vital role in the regulation of stem cells in human endometrium.

Another important finding of our study was that blocking Notch signaling appear to have the converse effect on the regeneration ability of mouse endometrial stem/progenitor cells. Using the BrdU pulse-chase approach, our group previously characterized LRSC from gestational and postpartum endometrium (23). In an artificial mouse endometrial breakdown and repair model, perivascular LRSC proliferated and contributed to remarkable stromal expansion during endometrial repair (38). Here in our established model, we observed similar findings. During decidualization, the number of LRSC decreased due to the dynamic changes in the endometrium. Although some LRSC were detected on the decidua tissue, a small population of non-proliferating LRSC were found in the basal region. These LRSC proliferated at breakdown, indicating endometrium can initiate repair at the time of shedding. Findings in human endometrium support this observation, whereby endometrial regeneration occurs simultaneously with tissue breakdown in a piecemeal fashion (39, 40). The proportion of LRSC pattern in our mouse menstrual-like model was similar to that reported by Kaitu’u-Lino *et al* (38).

Since it has been reported that Notch signaling has a regulatory role in uterine cellular remodeling throughout the oestrous cycle (41). It was not surprising that DAPT contribute to dysfunctional endometrial repair in the mouse menstrual-like model. Alternation in the stromal cells’ ability to proliferate may have contributed to size reduction of the repaired endometrium. Endometrial re-epithelialization is the first step in repair of the injured mucosal surfaces and this process was also delayed in DAPT treated mice. Before decidualization, ∼25.0% of Notch1^+^ LRSC were detected and the expression of Notch1 increased thereafter until late repair stage. This finding is consistent with a pregnant mouse model, whereby the initiation of implantation promoted expression of Notch1 in stromal cells (12). Interestingly, proliferating Notch1^+^ LRSC were only observed at decidualization, breakdown and early repair - the time course when the endometrium undergoes dynamic remodeling. These observations suggest the involvement of the Notch signaling during these events and more importantly the Notch-responsive stem cells are activated for tissue repair. Suppression of Notch signals in the uterine microenvironment significantly reduced the proliferation activity of Notch1^+^ LRSC and consequently the repair process was impaired. Similar results were observed in a Notch transcription factor *Rbpj* knockout mouse, where uterine-specific *Rbpj* knockout led to dysfunctional postpartum uterine repair (42).

In conclusion, our data demonstrate the importance of Notch signaling in regulating the activity of endometrial stem/progenitor cells *in vitro* and *in vivo*. Given its attenuation correlate with infertile women, understanding the precise mechanisms may provide a promising therapeutic approach in the near future.

## Materials and Methods

### Human tissues

Full thickness endometrial tissues were collected from 27 women aged 41– 52 years (mean age 46.4 years), who underwent abdominal hysterectomy for benign non-endometrial pathologies (Supplementary Table S1). Only pre-menopausal women with regular menstrual cycle and not taken any hormonal therapy for at least 3 months were recruited for this study. The phase of the endometrium was categorized as proliferative (n = 16) and secretory phase (n = 11) by an experienced histopathologist, who evaluated the hematoxylin eosin-stained endometrial sections of each sample. A written consent was signed by each patient after detailed counselling prior to participation of the study. Ethical approval was obtained from the Institutional Review Board of The University of Hong Kong/Hospital Authority Hong Kong West Cluster (UW20-465) and The Institutional Review Board of the University of Hong Kong-Shenzhen Hospital ([2018]94).

### Isolation of endometrial stromal cells

The isolation of single endometrial stromal cells was performed according to our previous study (16). In brief, endometrial tissue was minced into small pieces and digested with PBS containing collagenase type III (0.3 mg/ml, Worthington Biochemical Corporation, NJ, USA) and deoxyribonuclease type I (40 μg/ml, Worthington Biochemical Corporation) at 37°C for 1h. After two rounds of digestion, Ficoll-Paque (GE Healthcare, Uppsala, Sweden) centrifugation and anti-CD45 antibody coated Dynabeads (Invitrogen, Waltham, MA, USA) were sequentially used to remove the red blood cells and the leukocytes, respectively. Purified stromal cells were then separated from epithelial cells using anti-CD326 (EpCAM) antibody-coated microbeads (Miltenyi Biotec Inc., San Diego, CA, USA). Next, freshly isolated stromal cells were seeded into 100 mm dishes coated with fibronectin (1 mg/ml, Gibco) and cultured in growth medium (GM) containing 10% FBS (Invitrogen), 1% L-glutamine (Invitrogen) and 1% penicillin-streptomycin (Invitrogen) in DMEM/F-12 (Sigma-Aldrich, St Louis, MA, USA). Stromal cells were cultured in a humidified carbon dioxide incubator at 37°C. The medium was changed every 7 days until the cells reached 80% confluence.

### Magnetic bead selection for endometrial mesenchymal stem-like cells

EMSC were obtained by two sequential beadings with magnetic beads coated with anti-CD140b and anti-CD146 antibodies (16). Firstly, stromal cells were incubated with phycoerythrin (PE)-conjugated anti-CD140b antibody (R&D Systems, Minneapolis, MN, USA) for 45 mins at 4°C followed by another 15 mins incubation with anti-mouse IgG1 magnetic microbeads (Miltenyi Biotech). The obtained cell suspensions were then loaded onto MS columns (Miltenyi Biotech) with a magnetic field to separate the CD140b^+^ cells. The isolated CD140b^+^ stromal cells were cultured for 7-10 days to allow degradation of the microbeads during cell expansion. The cells were then trypsinized and incubated with anti-CD146 antibody coated microbeads (Miltenyi Biotec Inc.) for 15 mins at 4°C to obtain the CD140b^+^CD146^+^ cells for subsequent experiments. Phenotypic study of eMSC show their positive expression for CD140b and CD146 (16). Stromal cells at passage 1-3 were used in this study.

### Activation/inhibition of Notch or Wnt/β catenin signaling pathway

To activate Notch signaling, 48 well plates were coated with 50 µg/ml protein G (Invitrogen) overnight, washed with PBS and incubated with recombinant Jagged 1 protein (JAG1, 2 µg/ml, R&D System) for 3 h at room temperature. Freshly isolated eMSC (4 × 10^3^ cells/well) were seeded onto the JAG1 coated plates and cultured in GM for 7 days. To inhibit Notch signaling, eMSC were treated with N-[N-(3,5-difluorophenacetyl-l -alanyl)]-(S)-phenylglycine t-butyl ester (1.25 µM, DAPT, R&D Systems) and cultured in GM for 7 days.

For activation of WNT/β-catenin signaling, recombinant human WNT3A (0.01 μg/ml, R&D Systems) or recombinant human WNT5A (0.01 μg/ml, R&D Systems) was supplemented to eMSC cultured in GM. The WNT inhibitor, XAV939 (10 μM, R&D System) was used to inhibit the WNT signaling. GM containing DMSO was used as control.

### In vitro colony forming assay

Endometrial MSC were cultured on JAG1 coated plates for 7 days to form monolayer. These cells were then trypsinized, reseeded onto 6-well plates (400 cells/well) as passage 1 and cultured in GM for 14 days to form colonies. Individual colonies at passage 1 were harvested using cloning rings and the cells were reseeded onto 6-well plates (400 cells/well) for the subsequent passage and cultured for a further 14 days. Medium was changed every 7 days. The cloning efficiency was evaluated by the number of colonies divided by the number of seeded cells multiplied by 100.

### Cell proliferation assay

The CyQUANT™ NF Cell Proliferation kit (Thermo Scientific) was used to determine the proliferative ability of eMSC. In brief, eMSC (1 × 10^3^ cells/ well) were seeded into 96-well plates and cultured in GM. The cells were either treated with JAG1 or DAPT for 3 days. Cells treated with DMSO were used as control. After washing with PBS, 100 µl of dye binding solution was added to each well and cultured for 1 h at 37°C. Fluorescence intensity was measured by a fluorescence microplate reader with excitation at 485 nm and emission at 530 nm.

### Flow cytometry

Multi-color flow cytometry was applied to analyze the co-expression of CD140b and CD146 on endometrial stromal cells. The cells were incubated with PE-conjugated anti-CD140b antibody (2.5 μg/ml, PR7212 clone, Mouse IgG1, R&D Systems) and FITC-conjugated anti-CD146 antibodies (5 μg/ml, OJ79c clone, mouse IgG1; ThermoFisher Scientific) in dark for 45 mins at 4 °C. Fluorescent minus one (FMO) control was included for each antibody. Following the final washing step, the labeled cells were analyzed by a CytoFlex™ flow cytometer (Beckman Coulter, CA, USA). The cells were selected with electronic gating according to the forward and the side scatter profiles. Data were analyzed by the FlowJo Software (Tree Star Inc).

### G0/G1 cell cycle analysis

G0/G1 cell cycle analysis was performed as described (43). Endometrial MSC were treated with or without JAG1 or DAPT for 7 days. Then cells were trypsinized and incubated with APC-conjugated anti-CD140b antibody (2.5 μg/ml, PR7212 clone, Mouse IgG1, R&D Systems) and FITC-conjugated anti-CD146 antibodies (5 μg/ml, OJ79c clone, mouse IgG1; ThermoFisher Scientific) in dark for 45 mins at 4°C. The cells were then incubated with Hoechst dye (1 µg/ml) for 30 mins at 37°C. The Pyronin Y dye (1 μg/ml) was directly added to the cells and incubated for 30 mins at 37°C and analyzed using a Fortessa flow cytometer (BD Biosciences) at the Imaging and Flow cytometry Core, Centre for PanorOmic Sciences (CPOS), LKS Faculty of Medicine, The University of Hong Kong.

### WNT reporter assay

Endometrial MSC (2 – 5 × 10^4^ cells/well) were seeded into 24-well plate and cultured to 80% confluence when they were co-transfected with 4 μg of either TOP flash or FOP flash vector and 1 μg of pRL-TK (Renilla-TK-luciferase vector, Promega, Madison, WI, USA) using Lipofectamine 2000 (Invitrogen). Subsequently, the cells were treated with JAG1 or DAPT for 48 h. Some cells were treated 0.01µg/ml recombinant human WNT3A (R&D Systems) as positive control. The luciferase activity was measured using a GLOMAX™ 96 microplate luminometer. Transfection efficiency was determined by firefly luciferase activity normalized against the Renilla luciferase activity. The TOP/FOP ratio was presented as a measure of the TCF/LEF transcription.

### Fluorescence protein labeling assay

Human recombinant JAG1 (R&D System) was fluorescently labeled with Alexa Fluor™ 488 Microscale Protein Labeling Kit (Thermo Scientific) according to the manufacturer’s instructions. Briefly, JAG1 was incubated with 50 µL of sodium bicarbonate solution and Alexa Fluor dye for 1 h at room temperature. Then the protein-dye conjugate was flowed through a gel separation column (Bio-rad Biogel P-30) to purify the labeled protein, which was aliquoted and stored at -80°C for subsequent studies.

### Gene silencing

Endometrial MSC (2 × 10^4^/ well) were seeded onto 24-well plates, treated with JAG1 and cultured in OptiMEM medium (Invitrogen). After overnight incubation, the cells were transfected with 10 pmol of siRNA directed against Notch 1 (ID s9633 and s9635; Ambion) or random siRNA with scrambled sequence (Ambion) using the Lipofectamine RNAiMax transfection reagent (Invitrogen) according to the manufacturer’s instructions. The medium was changed to GM at 24 h post-transfection and the cells were trypsinized for flow cytometry analysis. The knockdown efficiency was determined by western blotting analysis (Fig S2F).

### Immunofluorescence staining of cells

For immunofluorescence staining, ∼8000 cells were cytospun at 12000 rpm for 10 mins and fixed with 4% paraformaldehyde for 10 mins. Permeabilization was conducted with 0.1% Triton-X 100 for 10 mins, followed by blocking with 5% BSA for 30 mins. Primary antibodies (Supplementary Table S2) or isotype-matched control antibodies were incubated overnight at 4°C. The next day, the corresponding secondary antibody (Supplementary Table S3) was added and incubated for 1 h. The cell nuclei were detected by staining with DAPI (Thermo Scientific) and mounted with mounting medium (Dako). The slides were washed with PBST between steps and all the steps were conducted at room temperature unless specified. Multi-spectrum fluorescence images were captured using a LSM 710 inverted confocal microscope and a LSM ZEN 2010 software (Carl Zeiss, Munich, Germany) at the CPOS, The University of Hong Kong. The total cell number and number of triple positive (CD140b+CD146+NICD+) cells were counted. At least 500 cells were counted from each sample.

### Western blot analysis

Cultured eMSC were lysed in cell lysis buffer (Ambion, Grandisland, NY, USA) in the presence of protease inhibitors. Then 5 μg of denatured protein samples were separated on 10% SDS-PAGE and transferred to polyvinylidene difluoride membranes (Immobilon™-P, Milllipore). After blocking with 5% skim milk for 1 h at room temperature, the membranes were incubated with appropriate primary antibodies overnight at 4°C followed by horseradish peroxidase conjugated secondary antibodies for 1h at room temperature (Supplementary Table S4). The protein expression was detected by the Western Bright ECL Kit (Advansta, CA, USA). The intensities of the western blot bands were quantified by the Quantity One software and normalized to that of β-actin.

### Quantitative real-time polymerase chain reaction

Quantitative real-time polymerase chain reaction (qPCR) with Taqman probes were used (Supplementary Table S5). The total RNA was isolated using the Absolutely RNA microprep kit (Agilent Technologies, Santa Clara, CA, USA) according to the manufacturer’s instructions. The concentration of total RNA was quantified by spectrophotometry. RNA was reversed transcribed to cDNA by the PrimeScript DNA Reverse Transcription kit (Takara, Japan). PCR was conducted by a 7500 Real-Time PCR System (Applied Biosystems). The mixtures were incubated at 50°C for 2 mins and 95 °C for 10 mins, followed by 40 cycles of 15s at 95°C and 1 min at 60°C. Gene expression was measured in triplicate and presented as relative gene expression using the 2^−ΔΔCt^ method and normalized to 18S as internal control.

### In situ proximity ligation assay (PLA)

The *in-situ* interaction of NICD and active β-catenin was determined using the Duolink™ II secondary antibodies and detection kits (Sigma–Aldrich, #DUO92001, #DUO92005 and #DUO92008) according to the manufacturer’s instructions. First, fixed cells were incubated with the PLA probes and primary antibodies against NICD and β-catenin (Supplementary Table S2) overnight at 4°C. The Duolink™ secondary antibodies were added in the following day and incubated at 37°C for 1h. The secondary antibodies were ligated together to form a closed circle by the Duolink™ ligation solution when the antibodies were in proximity (<40 nm) to each other. Polymerase and amplification buffer were then applied to amplify the positive signal (red dot) of the resulting closed circles and the cells were visualized with a LSM 800 inverted confocal microscope and a LSM ZEN 2010 software (Carl Zeiss, Munich, Germany) at the CPOS, The University of Hong Kong. The total number of cells and positive signals were counted. At least 500 cells were counted from each sample.

### Animal and housing condition

Mice were provided by Centre of Comparative Medicine Research at The University of Hong Kong. All experimental procedures performed in this study were approved by the Committee on Use of Live Animals in Teaching and Research, The University of Hong Kong, Hong Kong. The mice were kept under standard conditions with a light/dark cycle of 12h/12h and free access to food and water.

### Animal study design

The experimental setup is shown in Fig S6A. Day 19 prepubertal C57BL/6J female mice were labeled with BrdU according to our previous study (23). After a 6-week chase, the standard protocol to induce endometrial breakdown and repair was performed (22). In brief, female mice were mated with vasectomized >6-week-old C57BL/6J male mice (day 0). Pseudopregnant female mice were identified by the presence of a vaginal plug on the next day (day 1). On day 4 of pseudopregnancy, 30 µl of sesame oil was injected into the left uterine horn to induce decidualization while the right horn was not treated as control. The mice were euthanized, and their uteri were harvested on day 4 of pseudopregnancy (before decidualization), decidualization (day 7), breakdown (day 9), early repair (day 10) and late repair (day 12).

For the Notch inhibition study, decidualization was induced in both uterine horns on day 4 of pseudopregnancy. When endometrial breakdown occurred on day 9, DAPT (10 mg/kg) was injected into one uterine horn. The other uterine horn received the same volume of saline and served as a control. The uteri were collected on day 10 and day 12. The harvested tissues were fixed overnight with 4% paraformaldehyde and embedded in paraffin blocks for immunohistochemistry and immunofluorescent staining.

### Histological analysis of mouse endometrial thickness

Paraffin sections (5 µm) were stained with hematoxylin (Sigma-Aldrich) and eosin (Sigma-Aldrich) using standard protocols. Average endometrial thickness was measured from transverse section of the uterus - the vertical distance from the luminal epithelium to the endometrial–myometrial interface using the Image-Pro Plus software (version 6.0, Media Cybernetics) from 10 serial sections of the same animal.

### BrdU immunohistochemistry staining of mouse endometrial tissues

BrdU immunohistochemistry was performed as described (23). Paraffin sections were dewaxed and underwent antigen retrieval, followed by denaturation with 0.1 N HCl for 45 min. The sections were then quenched with 3% hydrogen peroxide for 10 min, blocked with 5% BSA/PBS for 1 h, and incubated with sheep anti-BrdU antibody (1:500 dilution; Abcam) or isotype control at 4°C overnight. On the next day, the sections were incubated with donkey anti-sheep biotinylated secondary antibodies (1:400 dilution; Abcam) for 1 h and then with the Vectastain ABC reagent (Vector Laboratories) for 30 min. BrdU positive staining was revealed by DAB solution (Dako) under a Zeiss Axioskop II microscope (Carl Zeiss). The sections were counterstained with the Mayer’s hematoxylin, dehydrated, and mounted using aqueous mounting medium (Dako). Images were acquired using a Photometrics CoolSNAP digital camera (Roper Scientific).

### Dual and triple immunofluorescence staining of mouse endometrial tissues

Paraffin sections underwent deparaffinization, rehydration, antigen denaturation and blocking as described above. For dual immunofluorescence staining, the two primary antibodies (Supplementary Table S2) were co-incubated at 4°C overnight. The slides were then incubated with the corresponding secondary antibodies (Supplementary Table S3) at 37°C for 1 h. For triple staining, the anti-BrdU antibody staining was conducted first, followed by incubation with the other two primary antibodies at 4°C overnight and subsequently the corresponding secondary antibodies on the next day. The slides were stained with DAPI (Thermo Scientific) and mounted with a fluorescence mounting medium (Dako). Multi-spectrum fluorescence images were captured by a Carl Zeiss LSM 800 inverted confocal microscope and the Zeiss LSM ZEN 2019 software (Carl Zeiss) at the CPOS, The University of Hong Kong.

### Enumeration of BrdU-Labeled Cells

BrdU-labeled cells were counted in a blinded manner as described (23). One transverse and one longitudinal section from each animal of different time points were analyzed. Images of the entire mouse uterine horn was acquired by a digital camera (Roper Scientific). The number of BrdU^+^ cells and the total number of cells in the stromal compartment were counted using the ImageJ software (NIH Image; National Institutes of Health). At least 2,000 nuclei per uterine horn per mouse at each time point were calculated. We only considered whole nuclei stained BrdU cells as LRSC. The percentage of BrdU^+^ cells were determined by dividing the number of BrdU^+^ cells by the total number of nuclei counted in each section.

### Statistical analysis

Data were analyzed using the GraphPad PRISM software (version 8.00; GraphPad Software Inc., San Diego, CA, USA). Distribution normality was tested using the Shapiro-Wilk test. Differences between two groups were analyzed using the Mann-Whitney U test for non-parametric data and the two-tailed unpaired Student’s t test for parametric data. Kruskal-Wallis test followed by Dunn’s post-test were used for multiple group comparison. Data are represented as mean ± SD. A difference with P-value of < 0.05 is considered as significant.

## Acknowledgments

We are grateful to all the women who agreed to donate their tissue samples for this study. We sincerely acknowledge Ms. Joyce Yuen, our project nurse and Miss Stella Wang, our research assistant. To all gynaecologists at Queen Mary Hospital and The University of Hong Kong Shenzhen Hospital for the collection of the samples. We are also grateful to the staffs at the Centre for PanorOmic Science (CPOS), Imaging and Flow cytometry Core and Centre of Comparative Medicine Research, The University of Hong Kong for their technical assistance in this study. This study was supported by funding from the National Natural Science Foundation of China/Research Grants Council Joint Research Scheme (N_HKU 732/20), the National Natural Science Foundation of China-Swedish Research Council Collaboration Research Programme (NSFC-VR 3181101648), the Shenzhen Knowledge Innovation Programme of the Shenzhen Science and Technology Innovation Commission (20180264) and The Hong Kong University Shenzhen Hospital Scientific Research Training Plan (HKUSZH20192003).

---

**Figure S1.**
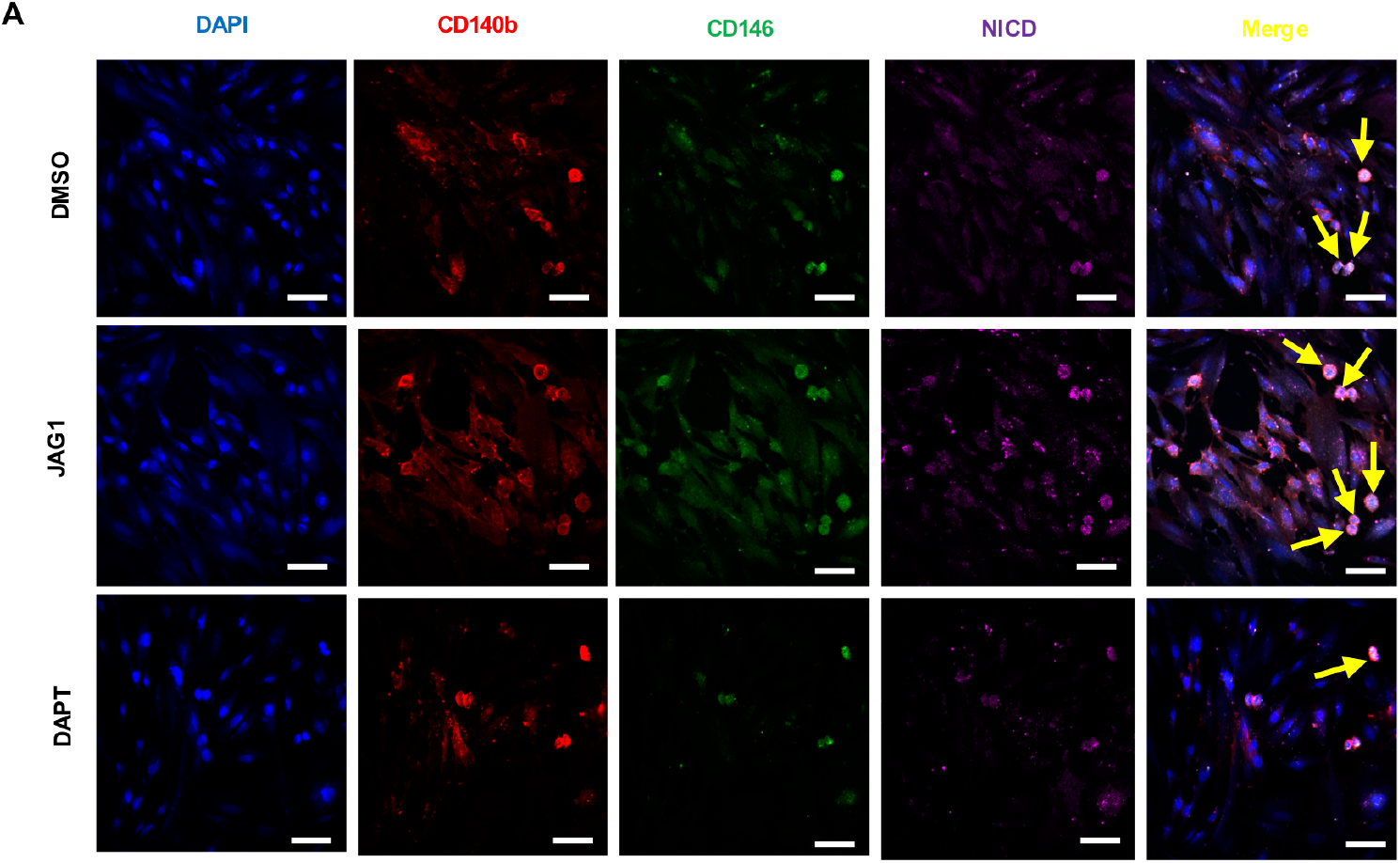
Representative images showing the triple staining of CD146+CD140b+NICD+ (yellow arrows) in endometrial stromal cells after Notch signals activation or inhibition. Scale bar: 50 µm, n=5.

**Figure S2.**
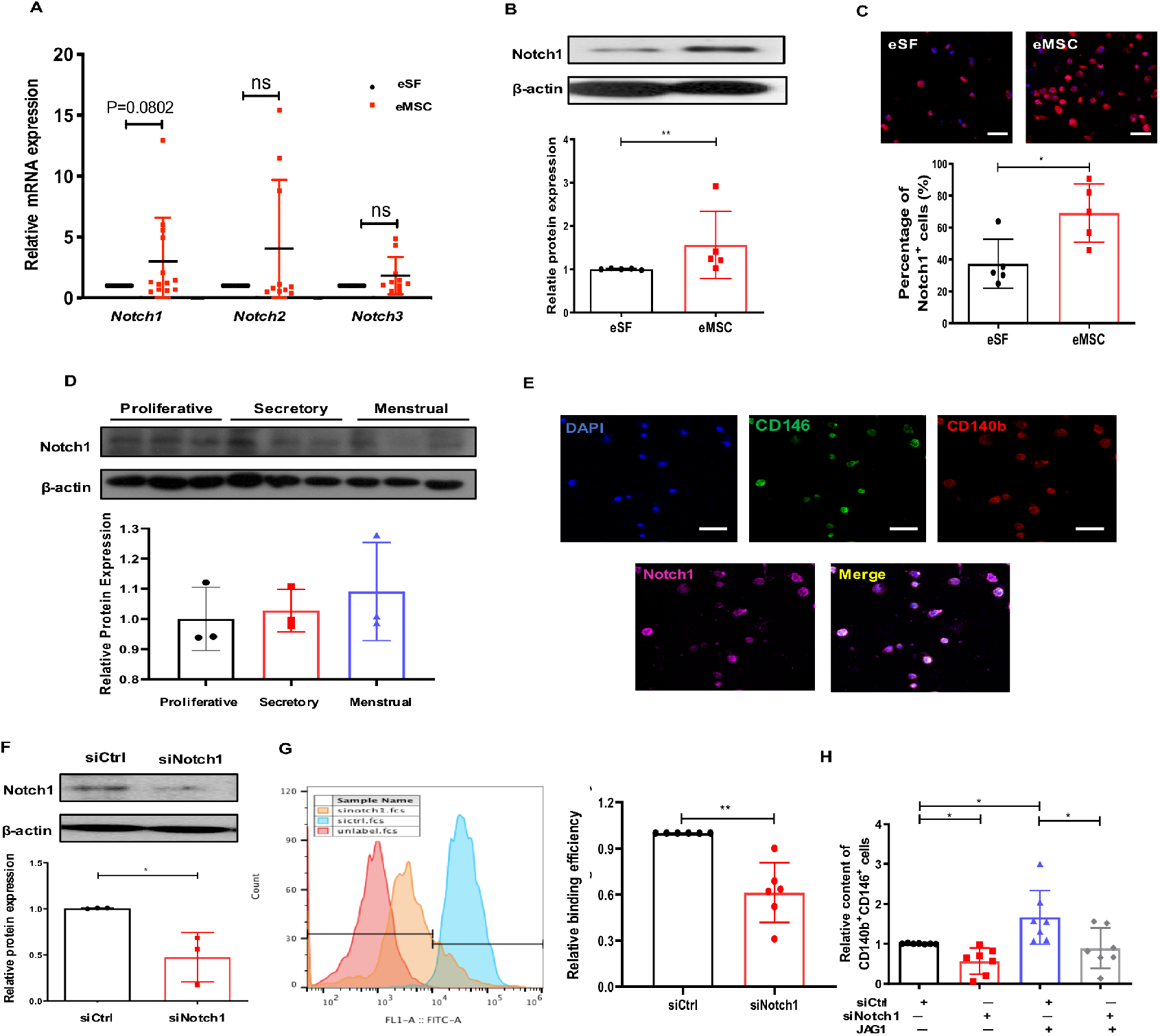
The role of Notch1 receptor on eMSC maintenance. **(A)** The relative gene expression of Notch receptor in eSF and eMSC (n=10-13). **(B)** Representative western blotting images and quantitative analysis of Notch1 protein expression in eSF and eMSC (n = 5). **(C)** Representative immunofluorescent images and quantitative analysis of Notch1 in eSF and eMSC (n=5), Scale bar: 50 µm. **(D)** Representative western blotting images and quantitative analysis of Notch1 protein expression in eMSC from different phase of menstrual cycle (n=3). **(E)** Representative immunofluorescent images showing the co-expression of CD140b (red), CD146 (green) and Notch1 (pink) on freshly isolated eMSC (n=5), scale bar: 50 µm. **(F)** Representative western blotting images and quantitative analysis of Notch1 protein expression on eMSC (n=3). **(G)** Representative images and quantitative analysis of binding efficiency between JAG1 and Notch 1 receptor on eMSC by flow cytometry (n=6). **(H)** The relative percentage of CD140b+CD146+ cells by flow cytometry (n=7). Results are presented as mean ± SD; *P < .05; **P < .01. Statistical analysis was performed using a two-tailed unpaired Student’s t test for parametric data and Mann-Whitney U test for non-parametric data, Kruskal-Wallis test followed by Dunn’s post-test for multiple group comparison.

**Figure S3.**
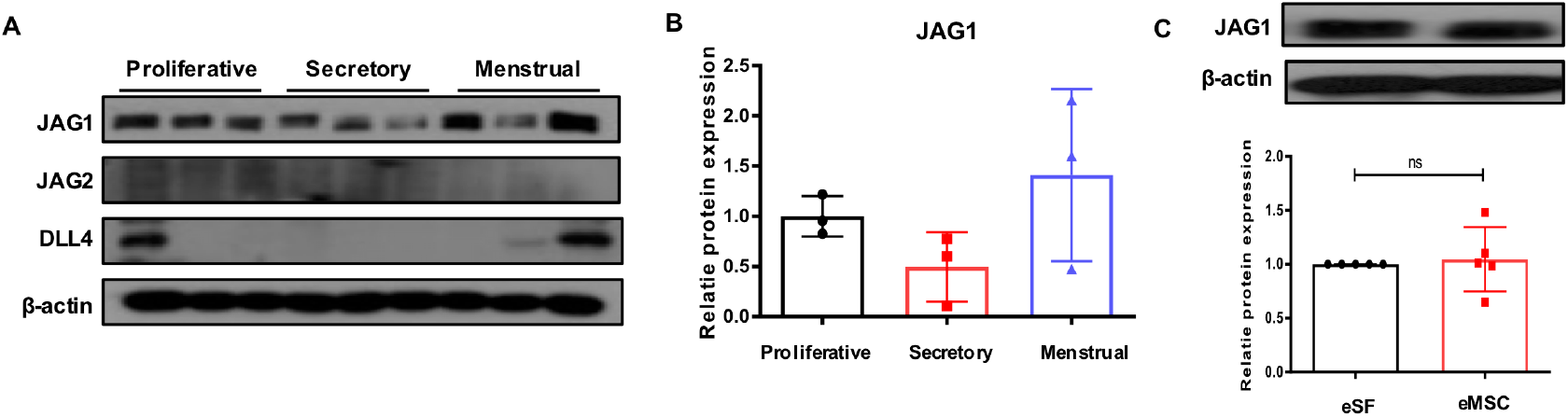
The expression of Notch ligands in eSF and eMSC. **(A)**Representative western blotting images of JAG1, JAG2, DLL4 in eMSC from different phase of the menstrual cycle. **(B)** Quantitative analysis of JAG1 protein expression (n=3). **(C)** Representative western blotting images and quantitative analysis of JAG1 protein expression in eSF and eMSC (n = 5). Statistical analysis was performed using a two-tailed unpaired Student’s t test for two group comparison, Kruskal-Wallis test followed by Dunn’s post-test for multiple group comparison. Abbreviation: eSF, endometrial stromal fibroblast; eMSC, endometrial mesenchymal stem-like cells.

**Figure S4.**
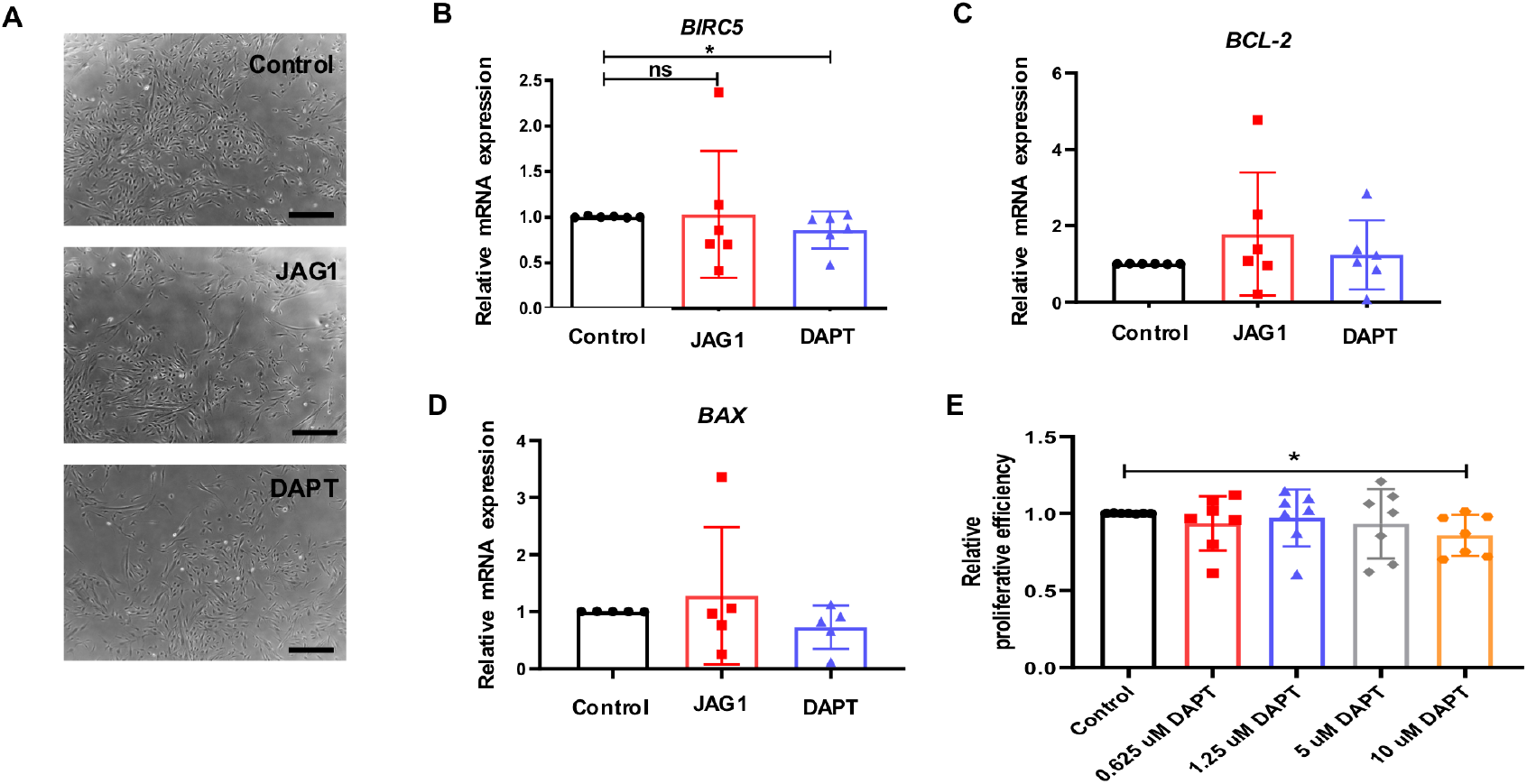
The effect of Notch signaling on eMSC apoptosis. **(A)** Representative bright field images of eMSC cultured under different condition for 7 days, scale bar: 100 µm. **(B-C)** The relative gene expression of anti-apoptosis marker *BIRC5* and *BCL-2* in eMSC after Notch signals activation or inhibition (n=5-6). **(D)** The relative gene expression of apoptosis marker *BAX* in eMSCs after Notch signals activation or inhibition (n=6). **(E)** The relative proliferative efficiency of eMSC under different DAPT concentration (n=7). Results are presented as mean ± SD. *P < .05; **P < .01. Statistical analysis was performed using Kruskal-Wallis test followed by Dunn’s post-test. Abbreviation: eSF, endometrial stromal fibroblast; eMSC, endometrial mesenchymal stem-like cells; rh, recombinant.

**Figure S5.**
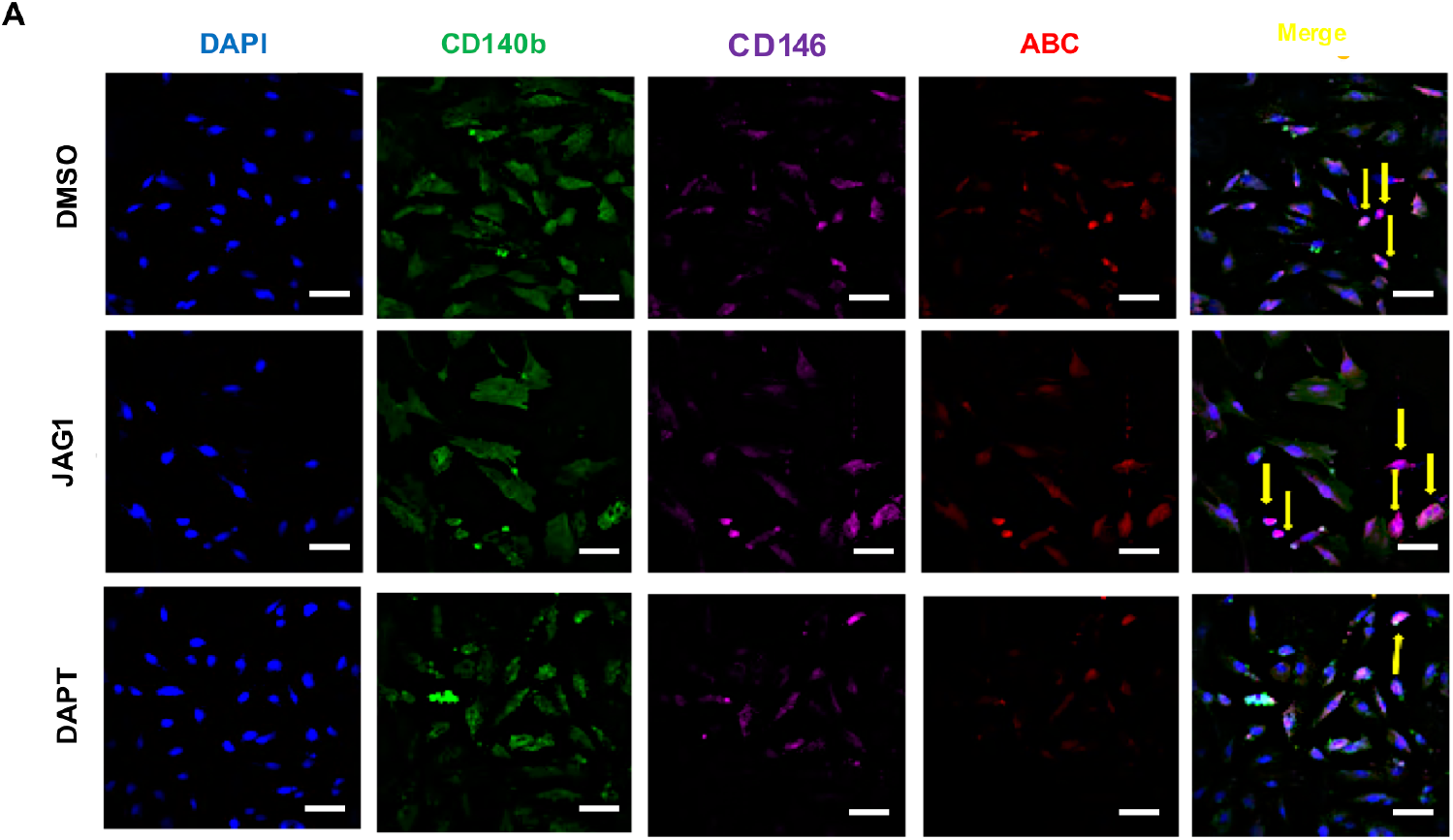
Representative images showing the triple staining of CD146+CD140b+ABC+ (yellow arrows) in endometrial stromal cells after Notch signals activation or inhibition. scale bar: 50 µm, n=5.

**Figure S6.**
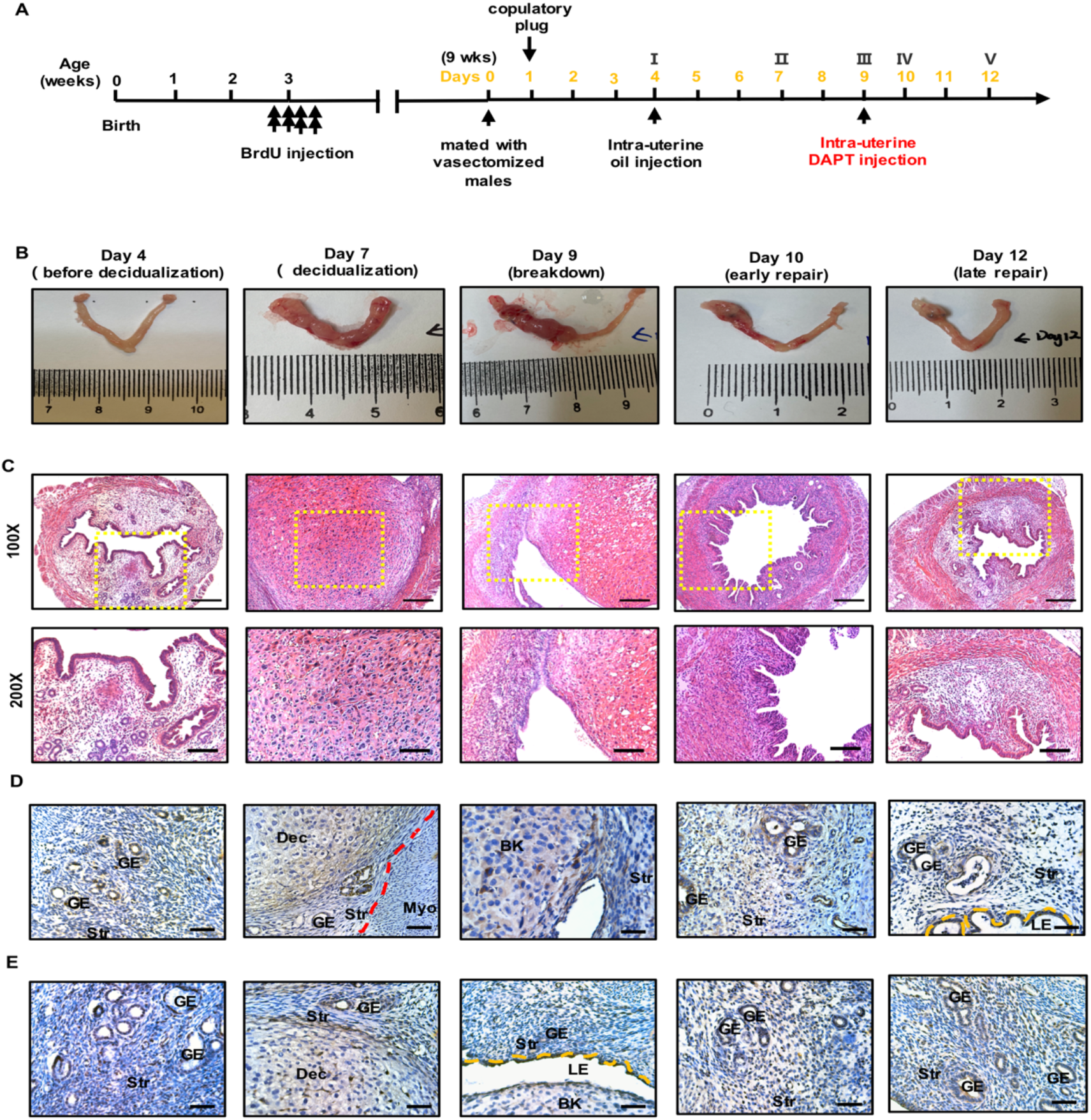
Establishment and histological analysis of mouse menstrual-like model. **(A)** Timeline of mouse menstrual-like model. **(B)** Macroscopic analysis of uteri at indicated time points. The right uterine horn severed as control. **(C)** Microscopic analysis of H&E-stained uteri at indicated time points. The lower panel are high magnification of images of the yellow square shown in the upper panel. Top panel, 100 µm; lower panel, 50 µm. **(D)** Representative immunohistology images of JAG1 on uterine tissues at different time points. Scale bar: 20 µm. **(E)** Representative immunohistology images of DLL4 on uterine tissues at different time points. Scale bar: 20 µm. n = 3–5 per group. Abbreviation: LE, luminal epithelium; GE, glandular epithelium; Str, stroma; Myo, myometrium; BD, breakdown; Dec, decidua.

**Figure S7.**
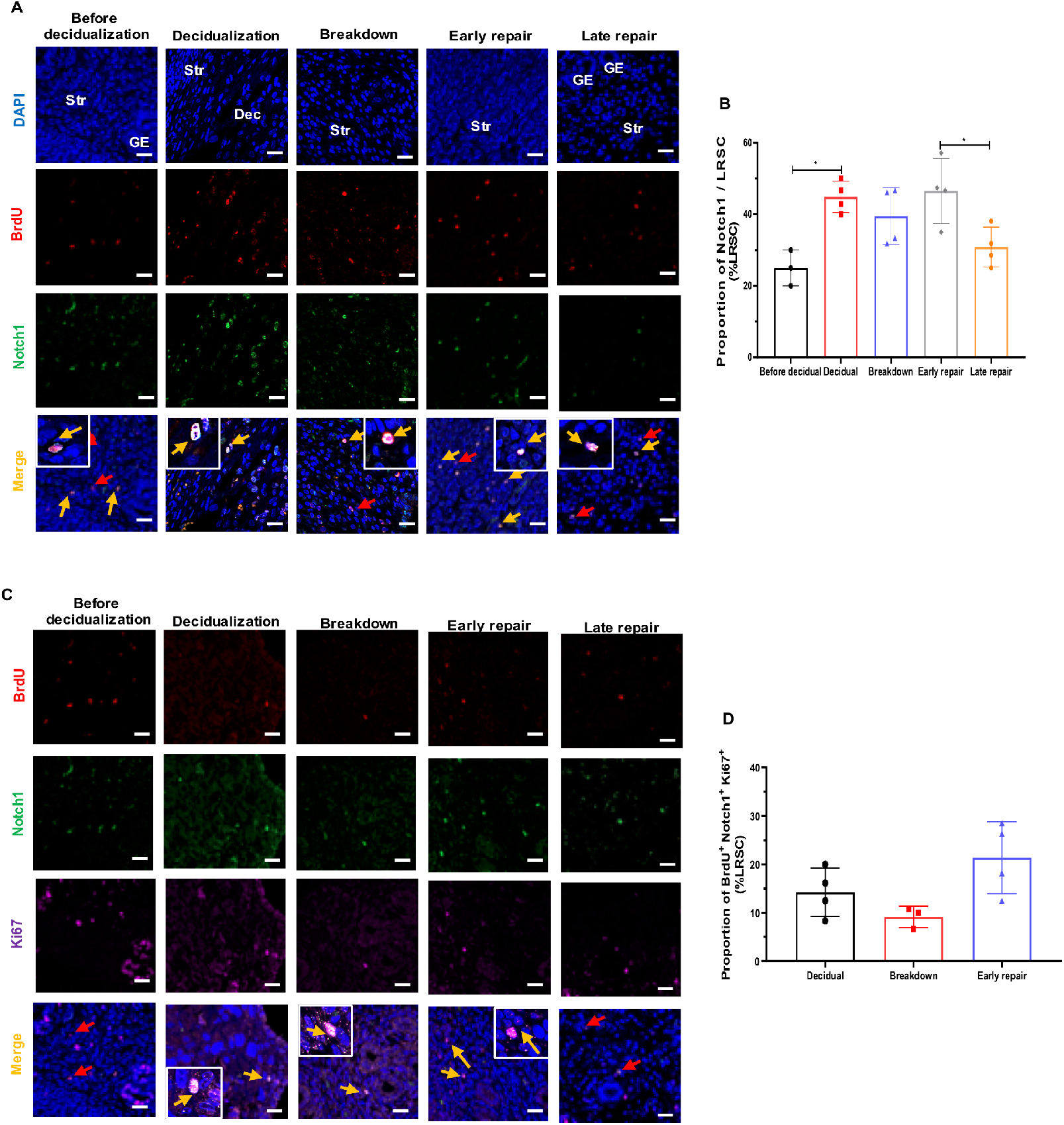
Characterization of LRSC in mouse menstrual-like model. **(A)** Representative immunofluorescent images show LRSC colocalizing with Notch1 (yellow arrow) before decidualization, decidualization, breakdown, early and late repair. Red arrows indicated no localization of Notch1 on LRSC. *Inserts* are enlarged figures of LRSC co-expressing Notch1. **(B)** The percentage of Notch1/ LRSC at different time points in mouse menstrual-like model. **(C)** Representative immunofluorescent images show proliferating LRSC colocalizing with Notch1(yellow arrow) at decidualization, breakdown and early repair. No colocalization with Notch1 (red arrow) for proliferating LRSC before decidualization and at late repair. Inserts are enlarged figures of LRSC co-expressing two proteins. **(D)** The percentage of BrdU^+^Notch1^+^Ki67^+^ at different time points in mouse menstrual-like model. Data are presented as mean ± SD; *P < .05. n = 3–5 per group. Scale bar: 20 µm. Statistical analysis was performed using a Kruskal-Wallis test followed by Dunn’s post-test. Abbreviation: BrdU, bromodeoxyuridine; Dec, decidual GE, glandular epithelium; Str, stroma.

**Figure S8.**
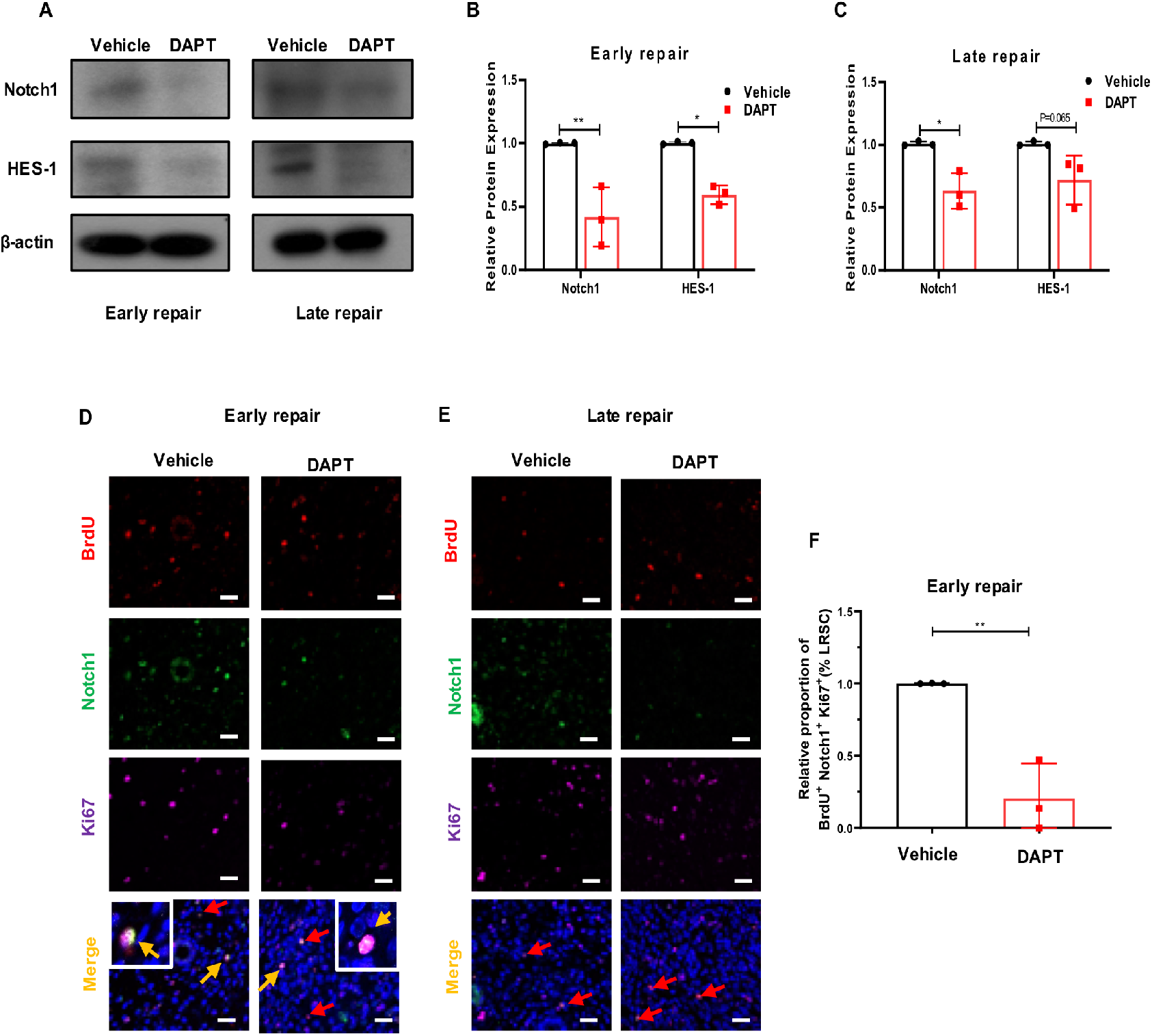
Characterization of LRSC after DAPT treatment in mouse menstrual-like model. **(A)** Representative western blotting images of Notch signals protein expression on mouse uterine tissue after DAPT treatment at different time points. **(B)** Quantitative analysis of Notch signals protein expression on mouse uterine tissue after DAPT treatment at early repair (n=3). **(C)** Quantitative analysis of Notch signals protein expression on mouse uterine tissue after DAPT treatment at late repair (n=3). **(D-E)** Representative immunofluorescent images show proliferating LRSC colocalizing with Notch1 (yellow arrow) after DAPT treatment at early repair and late repair. Red arrows indicated no localization of Notch1 on proliferating LRSC. Inserts are enlarged figures of LRSC co-expressing two proteins. **(F)** Relative percentage of BrdU^+^Notch1^+^Ki67^+^ cells after DAPT treatment at early repair. Data are presented as mean ± SD; *P < .05; **P < .01. n = 3–5 per group. Scale bar: 20 µm. Statistical analysis was performed using a two-tailed unpaired Student’s t test.

**Figure S9.**
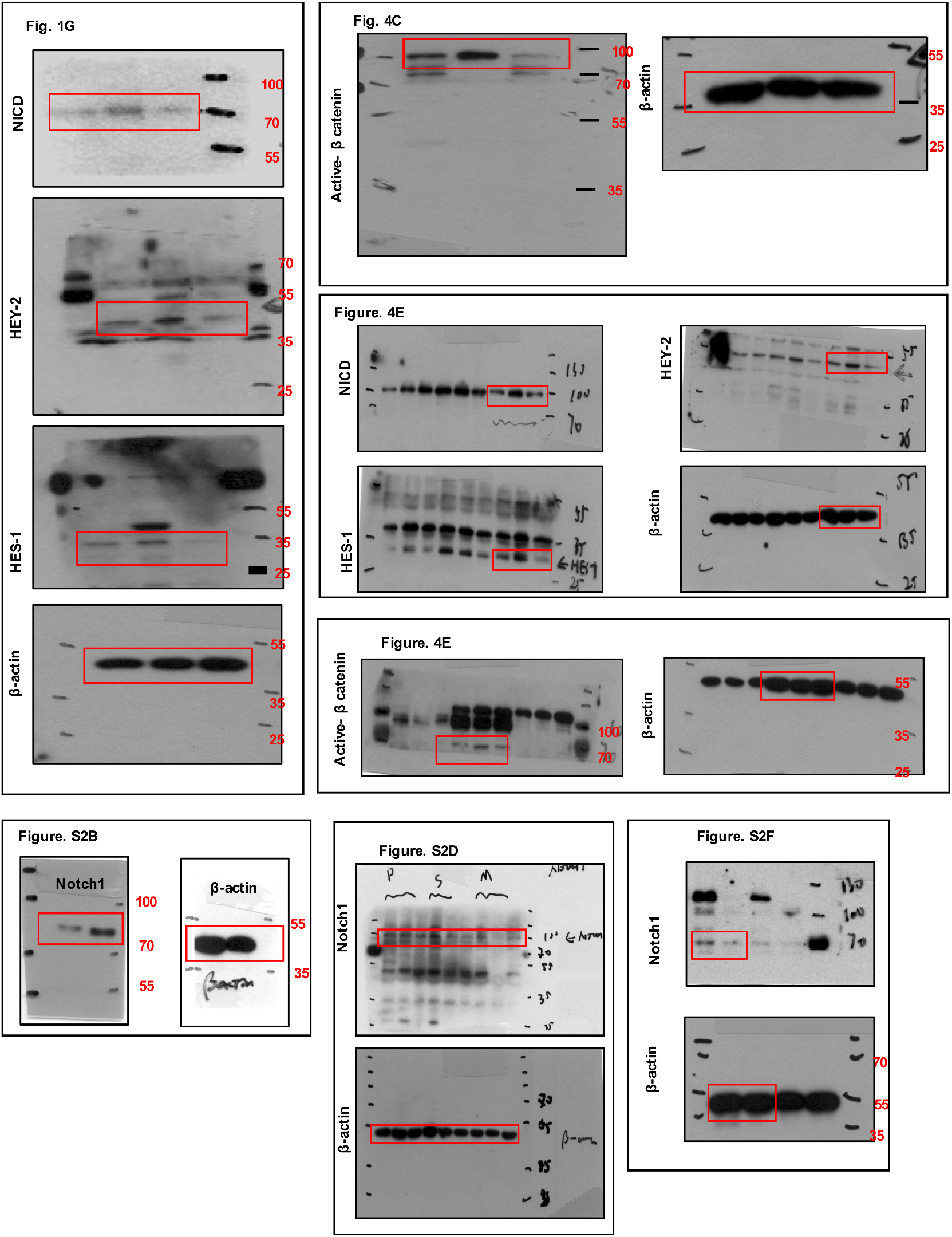
Uncropped scans of western blots. The cropped region is highlighted with the red boxes.

**Figure S10.**
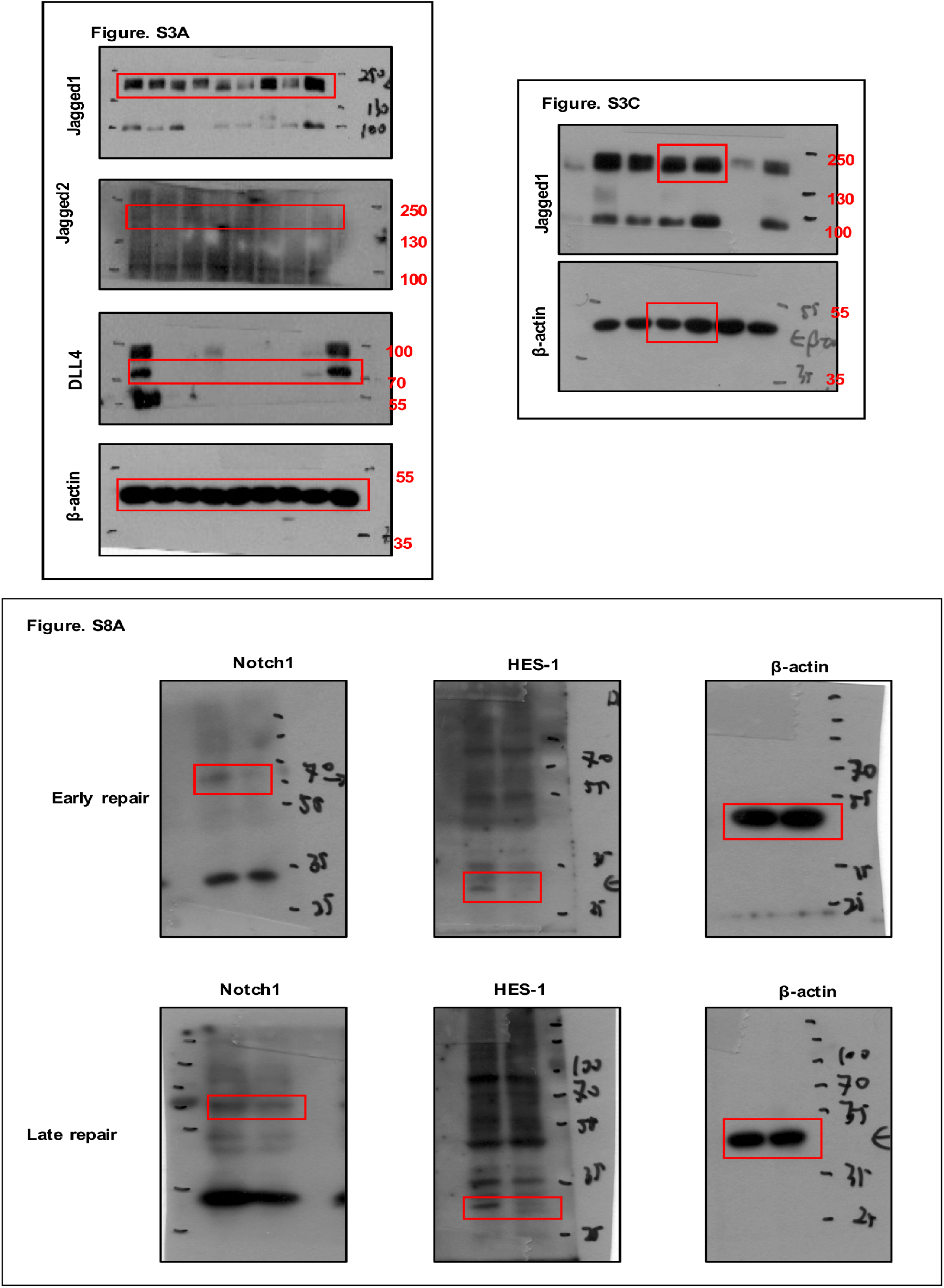
Uncropped scans of western blots. The cropped region is highlighted with the red boxes.

