## Supplementary file for "The Role of Notch Signaling in Endometrial Mesenchymal Stromal/Stem-like Cells Maintenance"

**Supplemental Table S1**. Pathological Characteristic of Full Thickness Endometrial

| | Age | Menstrual Phase | Pathology |
|---|---|---|---|
| 1 | 45 | proliferative | leiomyomas |
| 2 | 47 | proliferative | leiomyomas |
| 3 | 48 | proliferative | leiomyomas |
| 4 | 43 | proliferative | leiomyomas |
| 5 | 47 | proliferative | leiomyomas |
| 6 | 47 | proliferative | leiomyomas |
| 7 | 50 | proliferative | leiomyomas |
| 8 | 41 | proliferative | leiomyomas |
| 9 | 47 | proliferative | leiomyomas |
| 10 | 45 | proliferative | leiomyomas |
| 11 | 46 | proliferative | adenomyosis |
| 12 | 46 | proliferative | leiomyomas |
| 13 | 43 | proliferative | adenomyosis |
| 14 | 46 | proliferative | leiomyomas |
| 15 | 48 | proliferative | adenomyosis |
| 16 | 47 | proliferative | leiomyomas |
| 17 | 49 | secretory | adenomyosis |
| 18 | 52 | secretory | leiomyomas |
| 19 | 45 | secretory | leiomyomas |
| 20 | 44 | secretory | leiomyomas |
| 21 | 48 | secretory | leiomyomas |
| 22 | 44 | secretory | leiomyomas |
| 23 | 49 | secretory | leiomyomas |
| 24 | 44 | secretory | leiomyomas |
| 25 | 47 | secretory | leiomyomas |
| 26 | 46 | secretory | adenomyosis |
| 27 | 48 | secretory | leiomyomas |

**Supplemental Table S2**. List of primary antibodies used for immunohistochemistry (IHC) and immunofluorescent (IF) staining. Related to the results in Figure 4D, Figure 5, Figure6, Figure7, Figure S1, Figure S2, Figure S4, Figure S5, Figure S6 and Figure S7.

| Primary Antibody | Source | Dilution |
|---|---|---|
| **Active β-catenin:** mouse monoclonal to active β-actin | Millipore | 1:100 |
| **BrdU:** sheep polyclonal to BrdU | Abcam | 1:500 |
| **CD146:** mouse polyclonal to CD146 | Novus | 1:100 |
| **CD146:** rabbit polyclonal to CD146 | Abcam | 1:100 |
| **CD140b:** goat polyclonal to CD140b | Abcam | 1:100 |
| **Ki67:** rabbit polyclonal to ki-67 | Abcam | 1:500 |
| **Notch1:** rabbit polyclonal to Notch1 | Biorbyt | 1:100 |
| **Notch1 Intracellular domain (NICD):** rabbit polyclonal to NICD | Millipore | 1:100 |
| **Jagged1**: rabbit polyclonal to Jagged1 | CST | 1:200 |
| **DLL4:** rabbit polyclonal anti-DLL4 | Novus | 1:200 |

**Supplemental Table S3**. List of secondary antibodies used for immunohistochemistry (IHC) and immunofluorescent (IF) staining. Related to the results in Figure 4D, Figure 5, Figure6, Figure7, Figure S1, Figure S2, Figure S4, Figure S5, Figure S6 and Figure S7.

| Antibody | Conjugation | Source | Dilution |
|---|---|---|---|
| **Donkey anti-rabbit** | Alexa Fluor 647 | Life technologies | 1:200 |
| **Rabbit anti-mouse** | Alexa Fluor 488 | Life technologies | 1:200 |
| **Donkey anti-goat** | Alexa Fluor 555 | Life technologies | 1:200 |
| **Rabbit anti-goat** | Alexa Fluor 488 | Life technologies | 1:200 |
| **Donkey anti-mouse** | Alexa Fluor 568 | Life technologies | 1:200 |
| **Donkey anti-sheep** | Alexa Fluor 555 | Life technologies | 1:200 |
| **Donkey anti-sheep** | Biotin | Abcam | 1:400 |

**Supplemental Table S4**. List of primary and secondary antibodies used for western blotting. Related to the results in Figure 1, Figure 4, Figure S2 and Figure S7.

| Primary Antibody (Source, Dilution) | Secondary Antibody (Source, Dilution) |
|---|---|
| **Nocth1**: rabbit polyclonal to Nocth1 (Biorbyt, 1:500) | Rabbit horseradish peroxidase (GE Healthcare, 1:5000) |
| **Notch1 Intracellular domain (NICD)**: rabbit polyclonal to NICD (Millipore, 1:1000) | Rabbit horseradish peroxidase (GE Healthcare, 1:5000) |
| **Active β-catenin**: rabbit monoclonal to active β-actin (CST,1:1000) | Rabbit horseradish peroxidase (GE Healthcare, 1:5000) |
| **HEY-2**: rabbit polyclonal to HEY-2 (Millipore, 1:1000) | Rabbit horseradish peroxidase (GE Healthcare, 1:5000) |
| **HES-1**: rabbit polyclonal to HES-1 (Millipore, 1:500) | Rabbit horseradish peroxidase (GE Healthcare, 1:5000) |
| **Jagged1**: rabbit polyclonal to Jagged1 (CST, 1:1000) | Rabbit horseradish peroxidase (GE Healthcare, 1:5000) |
| **Jagged2**: rabbit polyclonal to Jagged2 (CST, 1:1000) | Rabbit horseradish peroxidase (GE Healthcare, 1:5000) |
| **DLL4**: rabbit polyclonal toDLL4 (CST, 1:1000) | Rabbit horseradish peroxidase (GE Healthcare, 1:5000) |
| **β-actin:** mouse monoclonal beta actin (Sigma, 1:5000) | Mouse horseradish peroxidase (GE Healthcare, 1:5000) |

**Supplemental Table S5.** Taqman probes used for qPCR. Related to RT- qPCR results in Figure 1, Figure 2, Figure S2A and Figure S3.

| **Gene Name *(Gene Symbol)*** | **Taqman Probes** |
|---|---|
| Hairy and enhancer of split-1 (*HES1)* | Hs00172878_m1 |
| Hairy/enhancer-of-split related with YRPW motif protein 1 *(HEY1)* | Hs01114113_m1 |
| Hairy/enhancer-of-split related with YRPW motif protein 2 *(HEY2)* | Hs01012057_m1 |
| Hairy/enhancer-of-split related with YRPW motif protein L *(HEYL)* | Hs01113778_m1 |
| Notch receptor 1 *(Notch1)* | Hs01062014_m1 |
| Notch receptor 2 *(Notch2)* | Hs01050702_m1 |
| Notch receptor 3 *(Notch3)* | Hs00166432_m1 |
| *CDKN1A* | Hs00355782_m1 |
| *CDKN1B* | Hs00153277_m1 |
| *CCNA2* | Hs00996788_m1 |
| *CCNE2* | Hs00180319_m1 |
| *CCND1* | Hs00765553_m1 |
| *GOS2* | Hs00377852_m1 |
| *MKI67* | Hs00757500_m1 |
| *BIRC5* | Hs04194392_m1 |
| *BAX* | Hs04986394_s1 |
| *BCL-2* | Hs04986394_s1 |
